# Escape from neutralizing antibodies by SARS-CoV-2 spike protein variants

**DOI:** 10.1101/2020.07.21.214759

**Authors:** Yiska Weisblum, Fabian Schmidt, Fengwen Zhang, Justin DaSilva, Daniel Poston, Julio C. C. Lorenzi, Frauke Muecksch, Magdalena Rutkowska, Hans-Heinrich Hoffmann, Eleftherios Michailidis, Christian Gaebler, Marianna Agudelo, Alice Cho, Zijun Wang, Anna Gazumyan, Melissa Cipolla, Larry Luchsinger, Christopher D. Hillyer, Marina Caskey, Davide F. Robbiani, Charles M. Rice, Michel C. Nussenzweig, Theodora Hatziioannou, Paul D. Bieniasz

**Affiliations:** Laboratory of Retrovirology, The Rockefeller University, 1230 York Avenue, New York NY 10065; Laboratory of Molecular Immunology The Rockefeller University, 1230 York Avenue, New York NY 10065; Laboratory of Virology and Infectious Disease The Rockefeller University, 1230 York Avenue, New York NY 10065; Lindsley F. Kimball Research Institute, New York Blood Center,New York, NY 10065; Institute for Research in Biomedicine, Università della Svizzera italiana, Bellinzona, Switzerland; Howard Hughes Medical Institute, The Rockefeller University, 1230 York Avenue, New York NY 10065

## Abstract

Neutralizing antibodies elicited by prior infection or vaccination are likely to be key for future protection of individuals and populations against SARS-CoV-2. Moreover, passively administered antibodies are among the most promising therapeutic and prophylactic anti-SARS-CoV-2 agents. However, the degree to which SARS-CoV-2 will adapt to evade neutralizing antibodies is unclear. Using a recombinant chimeric VSV/SARS-CoV-2 reporter virus, we show that functional SARS-CoV-2 S protein variants with mutations in the receptor binding domain (RBD) and N-terminal domain that confer resistance to monoclonal antibodies or convalescent plasma can be readily selected. Notably, SARS-CoV-2 S variants that resist commonly elicited neutralizing antibodies are now present at low frequencies in circulating SARS-CoV-2 populations. Finally, the emergence of antibody-resistant SARS-CoV-2 variants that might limit the therapeutic usefulness of monoclonal antibodies can be mitigated by the use of antibody combinations that target distinct neutralizing epitopes.

## Introduction

Neutralizing antibodies are a key component of adaptive immunity against many viruses that can be elicited by natural infection or vaccination (1). Antibodies can also be administered as recombinantly produced proteins or as convalescent plasma to confer a state of passive immunity in prophylactic or therapeutic settings. These paradigms are of particular importance given the emergence of SARS-CoV-2, and the devastating COVID19 pandemic that has ensued. Indeed, interventions to interrupt SAR-CoV-2 replication and spread are urgently sought, and passively administered antibodies are currently among the most promising therapeutic and prophylactic antiviral agents. Moreover, an understanding of the neutralizing antibody response to SARS-CoV-2 is critical for the elicitation of effective and durable immunity by vaccination (2).

Recent studies have shown that related, potently neutralizing monoclonal antibodies that recognize the SARS-CoV-2 receptor binding domain (RBD) are often elicited in SARS-CoV-2 infection (3-17). These antibodies have great potential to be clinically impactful in the treatment and prevention of SARS-CoV-2 infection. The low levels of somatic hypermutation and repetitive manner in which similar antibodies (e.g. those based on IGHV3-53 (3, 18, 19) have been isolated from COVID19 convalescents suggests that potently neutralizing responses should be readily elicited. Paradoxically, a significant fraction of COVID19 convalescents, including some from whom potent neutralizing antibodies have been cloned, exhibit low levels of plasma neutralizing activity (3, 20, 21). Together, these findings suggest that natural SARS-CoV-2 infection may often fail to induce sufficient B-cell expansion and maturation to generate high titer neutralizing antibodies.

The degree to, and pace at which SARS-CoV-2 might evolve to escape neutralizing antibodies is unclear. The aforementioned considerations raise the possibility that SARS-CoV-2 evolution might be influenced by frequent encounters with sub-optimal concentrations of potently neutralizing antibodies during natural infection. Moreover, the widespread use of convalescent plasma containing unknown, and often suboptimal, levels of neutralizing antibodies might also increase the acquisition of neutralizing antibody resistance by circulating SARS-CoV-2 populations (22, 23). Reinfection of previously infected individuals who have incomplete or waning serological immunity might similarly drive emergence of antibody escape variants. As human neutralizing antibodies are discovered and move into clinical development as therapeutics and prophylactics, and immunogens based on prototype SARS-CoV-2 spike protein sequences are deployed as vaccines, it is important to anticipate patterns of antibody resistance that might arise. Here, we describe a recombinant chimeric virus approach that can rapidly generate and evaluate SARS-CoV-2 S mutants that escape antibody neutralization. We show that mutations conferring resistance to convalescent plasma or RBD-specific monoclonal antibodies can be readily generated in vitro. Notably, these antibody resistance mutations are present at low frequency in natural populations. Importantly, the use of candidate monoclonal antibody combinations that target distinct epitopes on the RBD (and therefore have non-overlapping resistance mutations) can suppress the emergence of antibody resistance.

## Results

### Selection of SARS-CoV-2 S variants using a replication competent VSV/SARS-CoV-2 chimeric virus

To select SARS-CoV-2 S variants that escape neutralization by antibodies, we used a recently described replication-competent chimeric virus based on vesicular stomatitis virus that encodes the SARS-CoV-2 spike (S) protein and green fluorescent protein (rVSV/SARS-CoV-2/GFP) (24). Notably, rVSV/SARS-CoV-2/GFP replicates rapidly and to high titers (10^7^ to 10^8^ PFU/ml within 48h), mimics the SARS-CoV-2 requirement for ACE-2 as a receptor, and is neutralized by COVID19 convalescent plasma and SARS-CoV-2 S-specific human monoclonal antibodies (24). The replication of rVSV/SARS-CoV-2/GFP can be readily monitored and measured by GFP fluorescence and the absence of proof-reading activity in the viral polymerase (VSV-L) results in the generation of virus stocks with greater diversity than authentic SARS-CoV-2, for an equivalent viral population size. These features facilitate experiments to investigate the ability S protein variants to escape antibody neutralization.

We used two adapted, high-titer variants of rVSV/SARS-CoV-2/GFP, (namely rVSV/SARS-CoV-2/GFP_1D7_ and rVSV/SARS-CoV-2/GFP_2E1_) (24) in attempts to derive antibody-resistant mutants. Virus populations containing 1×10^6^ infectious particles were incubated with antibodies to neutralize susceptible variants (**Figure 1A**). For monoclonal antibodies, viral populations were incubated with antibodies at 5 μg/ml or 10μg/ml, (∼1000 to 10,000 x IC_50_). For plasma samples, viruses were incubated with a range of plasma dilutions (see materials and methods). Neutralized viral populations were then applied to 293T/ACE2(B) cells (24), which support robust rVSV/SARS-CoV-2/GFP replication and incubated for 48hr. We used three potent human monoclonal antibodies C121, C135 and C144 (3), that are candidates for clinical development (**Table 1**). In addition, we used four convalescent plasma samples, three of which were from the same donors from which C121, C135 and C144, were obtained (3) (**Table 1**).

**Table 1.** Plasma and monoclonal antibodies used in this study.

| Donor | Plasma NT<sub>50</sub><br>(rVSV-SARSCoV2/GFP) | Plasma NT<sub>50</sub><br>HIV/CCNGnLuc | Monoclonal<br>antibody |
| --- | --- | --- | --- |
| COV-47 | 6622 | 8016 | C144 |
| COV-72 | 6274 | 7982 | C135 |
| COV-107 | 122 | 334 | C121 |
| COV-NY | 12614 | 7505 | ND |

**Figure 1.**
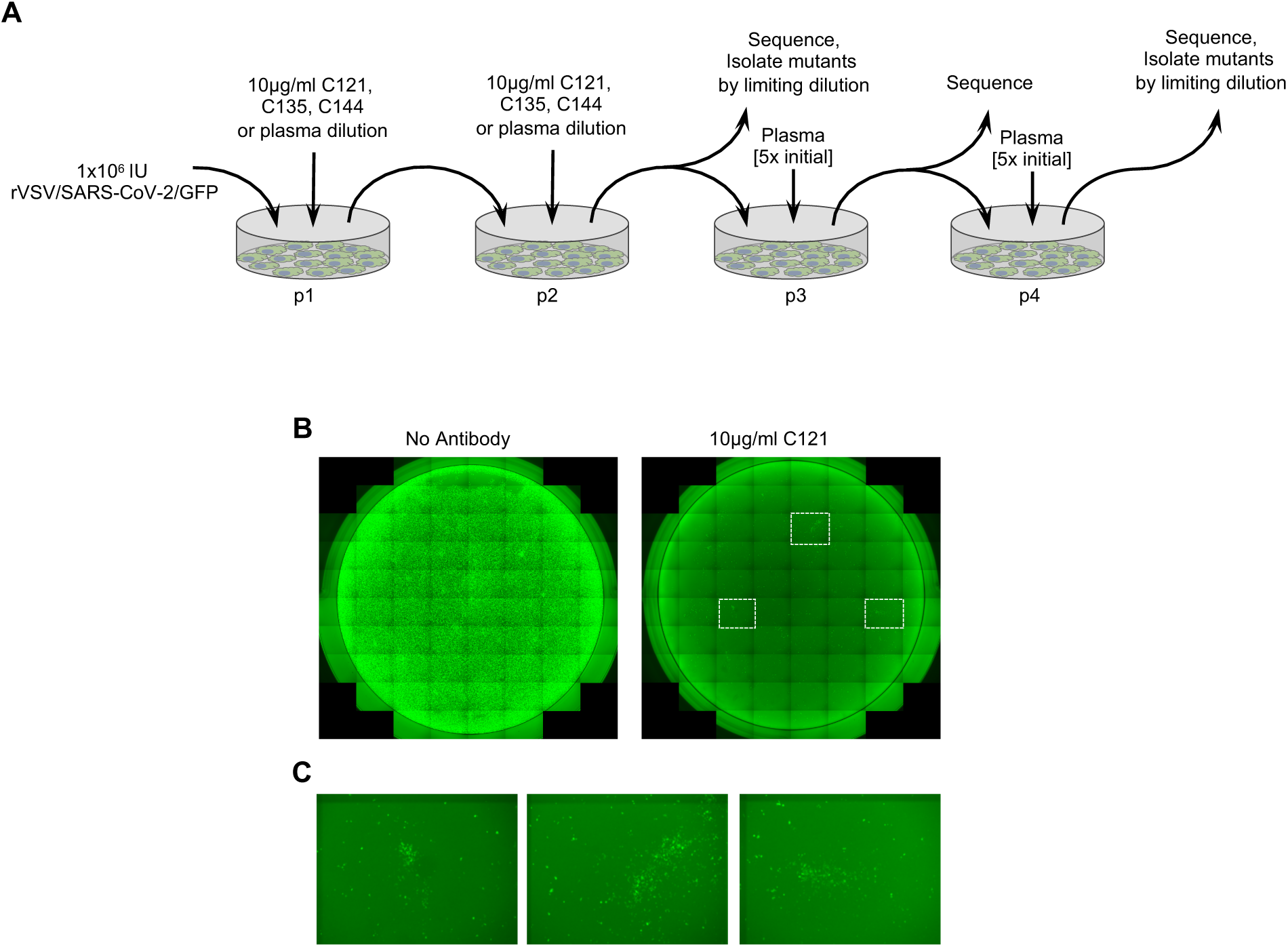
Selection of SARS-CoV-2 S mutations that confer antibody resistance. A. Outline of serial passage experiments with replication competent VSV derivatives encoding the SARS-CoV-2 S envelope glycoprotein and a GFP reporter (rVSV/SARS-CoV-2/GFP) in 293T/ACE2(B) cells in the presence of neutralizing antibodies or plasma. B. Representative images of 293T/ACE2(B) cells infected with 1×10^6^ PFU of rVSV/SARS-CoV-2/GFP in the presence or absence of 10μg/ml of the monoclonal antibody C121. C. Expanded view of the boxed areas showing individual plaques of putatively antibody resistant viruses.

Two of these plasmas (COV-47 and COV-72) were potently neutralizing while the third (COV-107) had low neutralizing activity. A fourth convalescent plasma sample (COV-NY) was potently neutralizing but did not have a corresponding monoclonal antibody (**Table 1**).

Infection with rVSV/SARS-CoV-2/GFP in the presence of the monoclonal antibodies C121 or C144 reduced the number of infectious units from 10^6^ to a few hundred, as estimated by the frequency of GFP-positive cells (**Figure 1B**) a reduction of >1,000 fold. C135 reduced infection by ∼100 fold. Imaging of wells infected with rVSV/SARS-CoV-2/GFP in the presence of C121 or C144 revealed a small number of foci (∼10 to 20/well), that suggested viral spread following initial infection (**Figure 1B**). In the case of C135, a greater number of GFP-positive cells were detected, obscuring the visualization of focal viral spread following initial infection. Aliquots of supernatants from these passage-1 (p1) cultures were collected 48h after infection, diluted in the same concentrations of monoclonal antibodies that were initially employed, and used to infect fresh (p2) cultures (**Figure 1A**). For p2 cultures, almost all cells became infected within 48h, suggesting the possible outgrowth of monoclonal antibody escape variants that were present in the original viral populations.

For selection in the presence of plasma, p1 supernatants were harvested at 48h after infection in the presence of the highest concentrations of plasma that permitted infection of reasonable numbers (approximately 10%) of cells. Then, p2 cultures were established using p1 supernatants, diluted in the same concentrations of plasma used in p1. This approach led to clear ‘escape’ for the COV-NY plasma with prolific viral growth in p2 as evidenced by a large increase in the number of GFP positive cells. For COV-47, COV-72 and CO107, plasma clearly retained at least some inhibitory activity in p2. Thereafter, p3 cultures and p4 cultures were established for COV-47, COV-72 and COV-107 plasmas at 5-fold higher concentrations of plasma than were used in p1 and p2 cultures (**Figure 1A**).

RNA was extracted from p2 supernatants (monoclonal antibodies and COV-NY plasma) as well as later passages for the COV-47, COV-72 and COV-107 plasma selections. Sequences encoding either the RBD or the complete S protein were amplified using PCR and analyzed by Sanger and/or Illumina sequencing. For all three monoclonal antibodies and two of the four plasmas, sequence analyses revealed clear evidence for selection, with similar or identical mutants emerging in the presence of monoclonal antibodies or plasma in both rVSV/SARS-CoV-2/GFP_1D7_ and rVSV/SARS-CoV-2/GFP_2E1_ cultures (**Figure 2A-D, Figure 2 - supplement 1A, B, Figure 2 - supplement 2A, B, Table 2**). In the case of C121, mutations E484K and Q493K/R within the RBD were present at high frequencies in both p2 selected populations, with mutation at a proximal position (F490L) present in one p2 population (**Figure 2A, Table 2)**. Viruses passaged in the presence of monoclonal antibody C144 also had mutations at positions E484 and Q493, but not at F490 (**Figure 2C, Table 2**). In contrast, virus populations passaged in the presence of monoclonal antibody C135 lacked mutations at E484 or Q493, and instead had mutations R346K/S/L and N440K at high frequency (**Figure 2C, Table 2**).

**Table 2.**
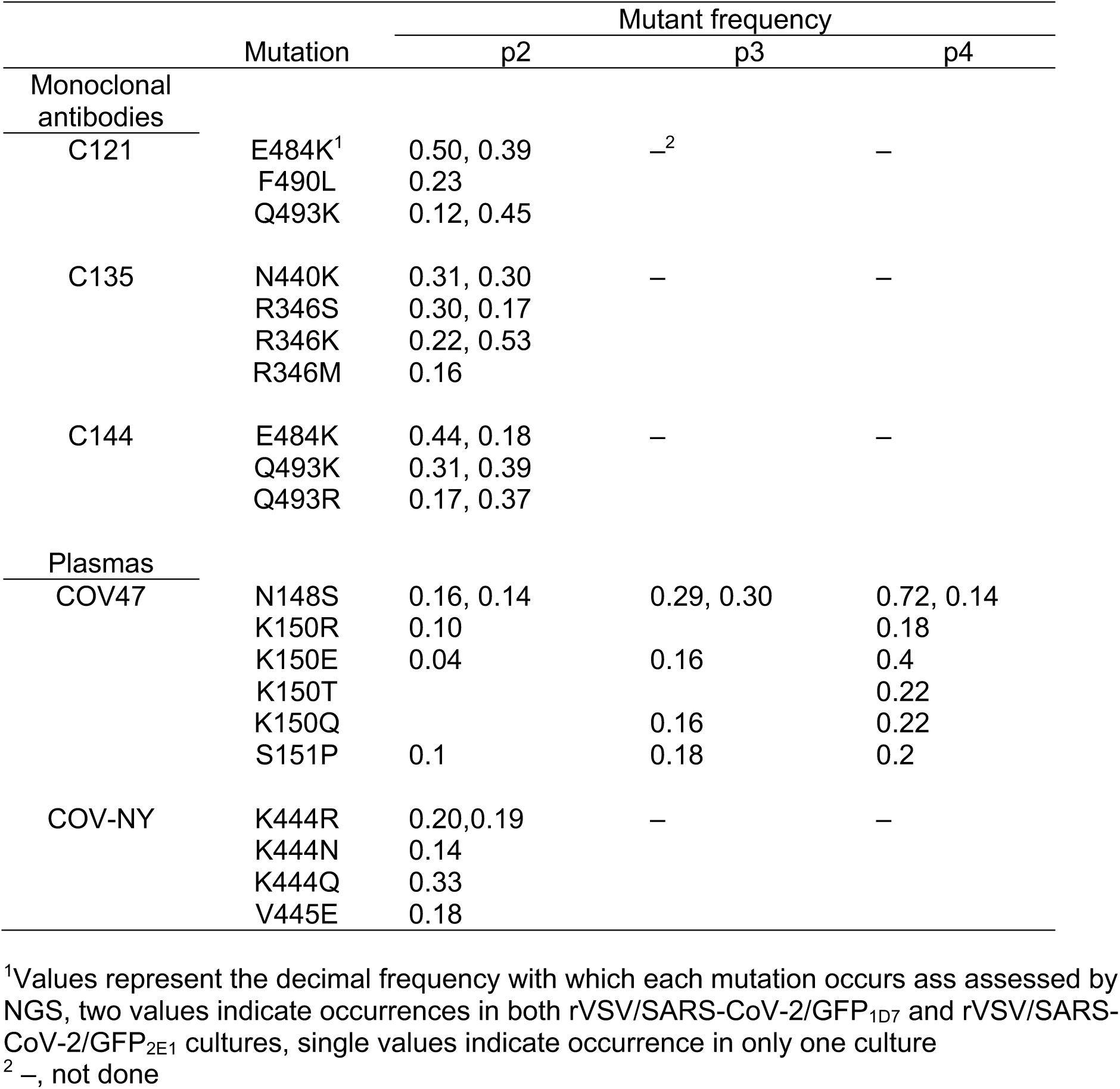
Mutations occurring at high frequency during rVSV/SARS-CoV-2 passage in the presence of neutralizing antibodies or plasma.

|  | Mutation | Mutant frequency |  |  |
| --- | --- | --- | --- | --- |
|  |  | p2 | p3 | p4 |
| Monoclonal antibodies |  |  |  |  |
| C121 | E484K<sup>1</sup> | 0.50, 0.39 | —<sup>2</sup> | — |
|  | F490L | 0.23 |  |  |
|  | Q493K | 0.12, 0.45 |  |  |
| C135 | N440K | 0.31, 0.30 | — | — |
|  | R346S | 0.30, 0.17 |  |  |
|  | R346K | 0.22, 0.53 |  |  |
|  | R346M | 0.16 |  |  |
| C144 | E484K | 0.44, 0.18 | — | — |
|  | Q493K | 0.31, 0.39 |  |  |
|  | Q493R | 0.17, 0.37 |  |  |
| Plasmas |  |  |  |  |
| COV47 | N148S | 0.16, 0.14 | 0.29, 0.30 | 0.72, 0.14 |
|  | K150R | 0.10 |  | 0.18 |
|  | K150E | 0.04 | 0.16 | 0.4 |
|  | K150T |  |  | 0.22 |
|  | K150Q |  | 0.16 | 0.22 |
|  | S151P | 0.1 | 0.18 | 0.2 |
| COV-NY | K444R | 0.20, 0.19 | — | — |
|  | K444N | 0.14 |  |  |
|  | K444Q | 0.33 |  |  |
|  | V445E | 0.18 |  |  |

<sup>1</sup>Values represent the decimal frequency with which each mutation occurs as assessed by NGS, two values indicate occurrences in both rVSV/SARS-CoV-2/GFP<sub>1D7</sub> and rVSV/SARS-CoV-2/GFP<sub>2E1</sub> cultures, single values indicate occurrence in only one culture
<sup>2</sup> —, not done

**Figure 2.**
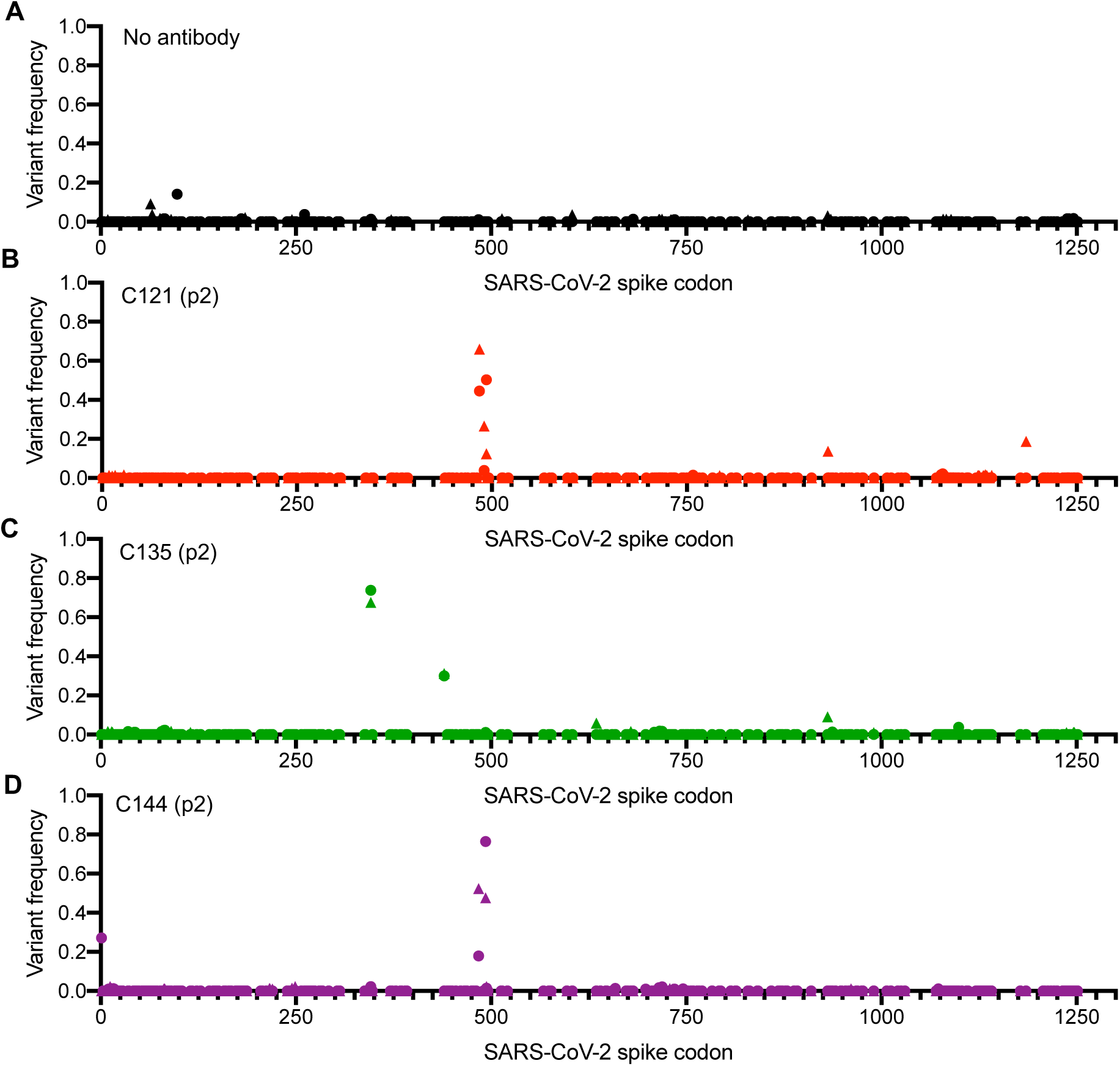
Analysis of S-encoding sequences following rVSV/SARS-CoV-2/GFP replication in the presence of neutralizing monoclonal antibodies. A-D. Graphs depict the S codon position (X-axis) and the frequency of non-synonymous substitutions (Y-axis) following the second passage (p2) of rVSV/SARS-CoV-2/GFP on 293T/ACE2(B) cells in the absence of antibody or plasma (A), or in the presence of 10μg/ml C121 (B), C135 (C) or C144 (D). Results are shown for both rVSV/SARS-CoV-2/GFP variants (the frequency of 1D7 mutations is plotted as circles and 2E1 mutations as triangles).

Mutations at specific positions were enriched in viruses passaged in the presence of convalescent plasma, in 2 out of 4 cases (**Figure 2 - supplement 1A, B, Figure 2 - supplement 2A, B, Table 2**). Specifically, virus populations passaged in the presence of COV-NY plasma had mutations within RBD encoding sequence (K444R/N/Q and V445E) that were abundant at p2 (**Figure 2 - supplement 2A, B, Table 2**). Conversely, mutations outside the RBD, specifically at N148S, K150R/E/T/Q and S151P in the N-terminal domain (NTD) were present at modest frequency in COV-47 p2 cultures and became more abundant at p3 and p4 (**Figure 2 - supplement 1A, B, Table 2**). Replication in the presence of COV72 or COV107 plasma did not lead to the clear emergence of escape mutations, suggesting that the neutralization by these plasmas was not due to one dominant antibody specificity. In the case of COV107, the failure of escape mutants to emerge may simply be due to the lack of potency of that plasma (**Table 1**). However, in the case of COV-72, combinations of antibodies may be responsible for the potent neutralizing properties of the plasma in that case.

### Isolation and characterization of rVSV/SARS-CoV-2/GFP antibody escape mutants

Based on the aforementioned analyses, supernatants from C121, C144, and C135 and COV-NY plasma p2 cultures, or COV47 p4 cultures, contained mixtures of putative rVSV/SARS-CoV-2/GFP neutralization escape mutants. To isolate individual mutants, the supernatants were serially diluted and individual viral foci isolated by limiting dilution in 96 well plates. Numerous individual rVSV/SARS-CoV-2/GFP_1D7_ and rVSV/SARS-CoV-2/GFP_2E1_ derivatives were harvested from wells containing a single virus plaque, expanded on 293T/ACE2(B) cells, then RNA was extracted and subjected to Sanger sequencing (**Figure 3 - supplement 1**). This process verified the purity of the individual rVSV/SARS-CoV-2/GFP variants and yielded a number of viral mutants for further analysis (**Figure 3 - supplement 1**). These mutants all encoded single amino-acid substitutions in S-coding sequences that corresponded to variants found at varying frequencies (determined by Illumina sequencing) in the parental viral populations. Notably, each of the isolated rVSV/SARS-CoV-2/GFP mutants replicated with similar kinetics to the parental rVSV/SARS-CoV-2/GFP_1D7_ and rVSV/SARS-CoV-2/GFP_2E1_ viruses (**Figure 3A**), suggesting that the mutations that emerged during replication in the presence of monoclonal antibodies or plasma did not confer a substantial loss of fitness, at least in the context of rVSV/SARS-CoV-2/GFP. Moreover, for mutants in RBD sequences that arose in the C121, C135, C144 and COV-NY cultures, each of the viral mutants retained approximately equivalent sensitivity to neutralization by an ACE2-Fc fusion protein, suggesting little or no change in interaction with ACE2 (**Figure 3B**).

**Figure 3.**
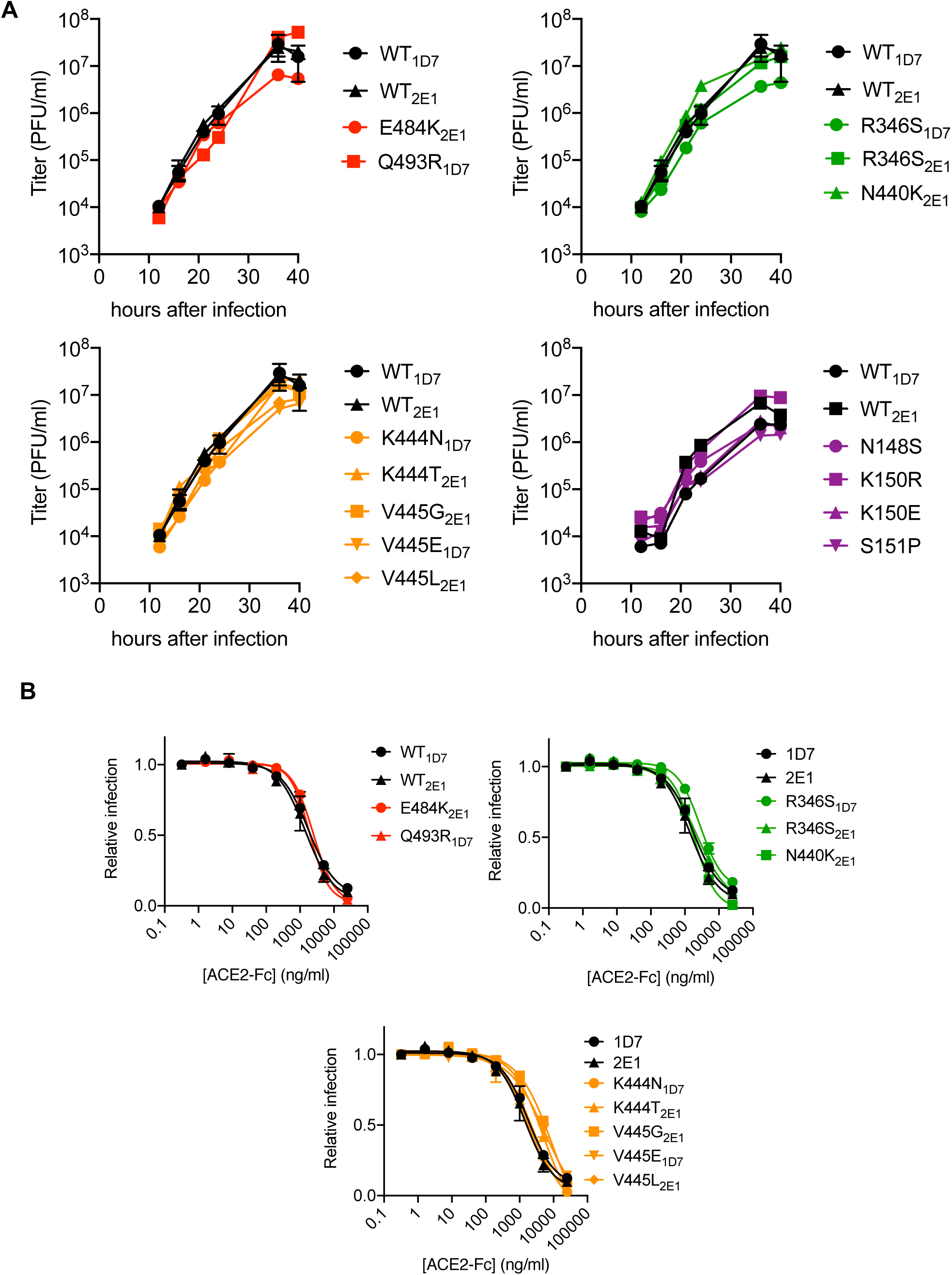
Characterization of mutant rVSV/SARS-CoV-2/GFP derivatives. A. Replication of plaque purified rVSV/SARS-CoV-2/GFP bearing individual S amino acid substitutions that arose during passage with the indicated antibody or plasma. 293T/ACE2cl.22 cells were inoculated with equivalent doses of parental or mutant rVSV/SARS-CoV-2/GFP isolates. Supernatant was collected at the indicated times after inoculation and number of infectious units present therein was determined on 293T/ACE2cl.22 cells. The mean of two independent experiments is plotted. B. Infection 293T/ACE2cl.22 cells by rVSV/SARS-CoV-2/GFP encoding the indicated S protein mutations in the presence of increasing amounts of a chimeric ACE2-Fc molecule. Infection was quantified by FACS. Mean of two independent experiments.

We next determined the sensitivity of the isolated RBD mutants to neutralization by the three monoclonal antibodies. The E484K and Q493R mutants that emerged during replication in the presence of C121 or C144, both caused apparently complete, or near complete, resistance to both antibodies (IC_50_ >10 μg/ml, **Figure 4A, B**). However, both of these mutants retained full sensitivity (IC_50_ <10 ng/ml) to C135. Conversely, the R346S and N440K mutants that emerged during replication in the presence of C135 were resistant to C135, but retained full sensitivity to both C121 and C144 (**Figure 4A, B**). The K444N, K444T, V445G, V445E and V445L mutants that arose during replication the presence of COV-NY plasma conferred partial resistance to C135, with IC_50_ values ranging from 25ng/ml to 700ng/ml, but these mutants retained full sensitivity to both C121 and C144 (**Figure 4A, B**). The spatial distribution of these resistance-conferring mutations on the SARS-CoV-2 S RBD surface suggested the existence of both distinct and partly overlapping neutralizing epitopes on the RBD (**Figure 4C**). The C121 and C144 neutralizing epitopes appear to be similar, and include E484 and Q493, while a clearly distinct conformational epitope seems to be recognized by C135, that includes R346 and N440 residues. Antibodies that constitute at last part of the neutralizing activity evident in COV-NY plasma appear to recognize an epitope that includes and K444 and V445. The ability of mutations at these residues to confer partial resistance to C135 is consistent with their spatial proximity to the C135 conformational epitope (**Figure 4C**).

**Fig. 4.**
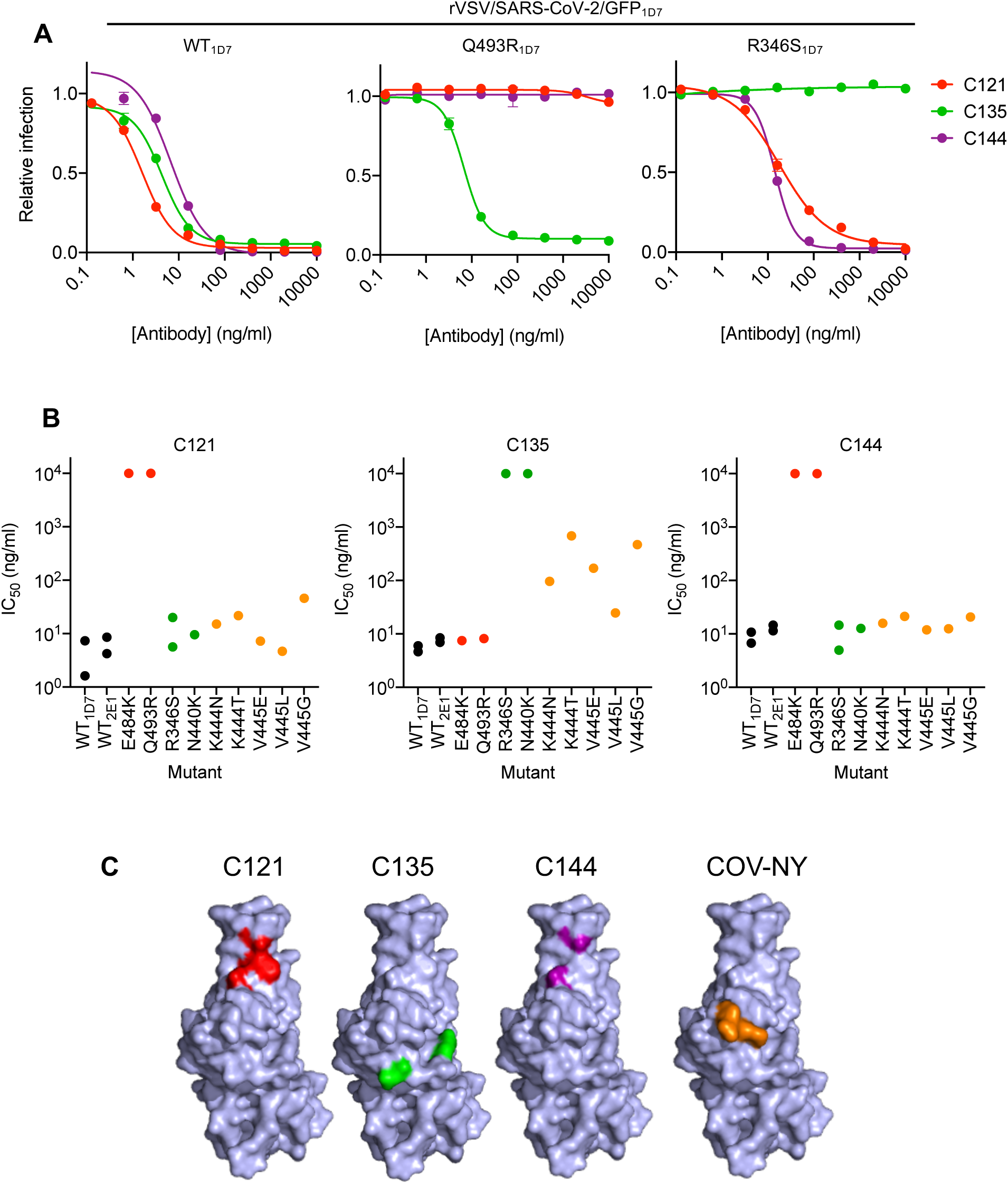
Neutralization of rVSV/SARS-CoV-2/GFP RBD mutants by monoclonal antibodies. A. Examples of neutralization resistance of rVSV/SARS-CoV-2/GFP mutants that were isolated following passage in the presence of antibodies. 293T/ACE2cl.22 cells were inoculated with WT or mutant rVSV/SARS-CoV-2/GFP in the presence of increasing amount of each monoclonal antibody, and infection quantified by FACS 16h later. Mean and SD from two independent experiments. B. Neutralization sensitivity/resistance of rVSV/SARS-CoV-2/GFP mutants isolated following replication in the presence of antibodies. IC_50_ values were calculated for each virus-monoclonal antibody combination in two independent experiments. C. Position of neutralization resistance-conferring substitutions. Structure of the RBD (from PDB 6M17 (32)) with positions that are occupied by amino acids where mutations were acquired during replication in the presence of each monoclonal antibody or COV-NY plasma indicated.

To test whether neutralization escape mutations conferred loss of binding to the monoclonal antibodies, we expressed conformationally prefusion-stabilized S trimers (25), appended at their C-termini with NanoLuc luciferase (**Figure 5A**). The S-trimers were incubated in solution with the monoclonal antibodies, complexes were captured using protein G magnetic beads, and the amount of S-trimer captured was measured using NanoLuc luciferase assays (**Figure 5A**). As expected, C121, C135 and C144 monoclonal antibodies all bound the WT S trimer (**Figure 5B**). The E484K and Q493R trimers exhibited complete, or near complete loss of binding to C121 and C144 antibodies but retained WT levels of binding to C135 (**Figure 5B**). Conversely, the R346S and N440K mutants exhibited complete loss of binding to C135, but retained WT levels of binding to C121 and C144. The K444N and V445E mutants retained near WT levels of binding to all three antibodies, despite exhibiting partial resistance to C135 (**Figure 5A, B**). Presumably the loss of affinity of these mutants for C135 was sufficient to impart partial neutralization resistance but insufficient to abolish binding in the solution binding assay.

**Fig. 5.**
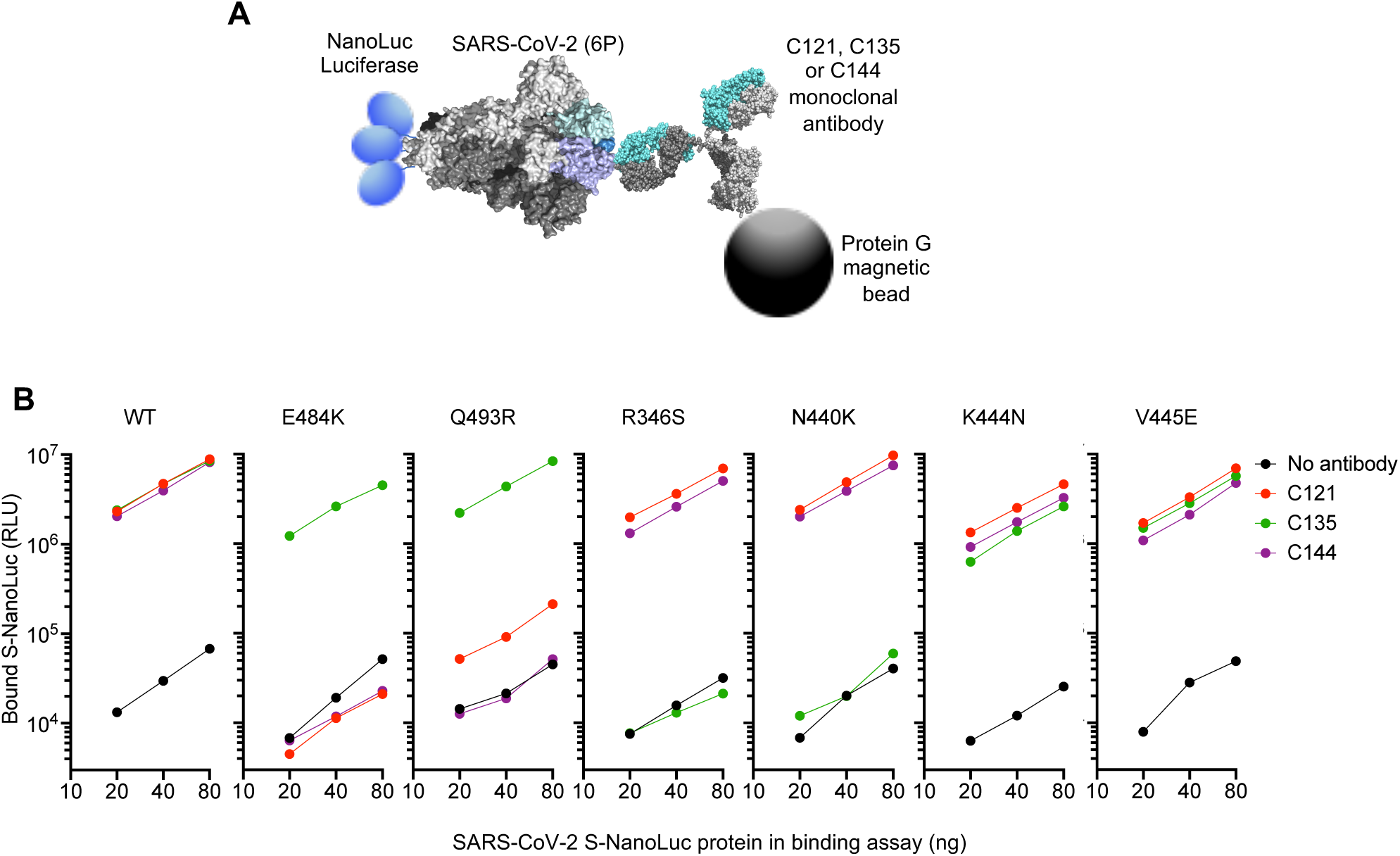
Loss of binding to monoclonal antibodies by neutralization escape mutants. A. Schematic representation of the binding assay in which NanoLuc luciferase is appended to the C termini of a conformationally stabilized S-trimer. The fusion protein is incubated with antibodies and complexes captured using protein G magnetic beads B. Bound Nanoluc luciferase quantified following incubation of the indicated WT or mutant Nanoluc-S fusion proteins with the indicated antibodies and Protein G magnetic beads.

Analysis of mutants that were isolated from the virus population that emerged during rVSV/SARS-CoV-2/GFP replication in the presence of COV-47 plasma (specifically N148S, K150R, K150E, S151P) revealed that these mutants exhibited specific resistance to COV-47 plasma. Indeed, the COV-47 plasma NT_50_ for these mutants was reduced by 8-to 10-fold (**Figure 6A**). This finding indicates that the antibody or antibodies responsible for majority of neutralizing activity in COV-47 plasma target an NTD epitope that includes amino acids 148-151, even though the highly potent monoclonal antibody (C144) isolated from COV-47 targets the RBD. Mutants in the 148-151 NTD epitope exhibited marginal reductions in sensitivity to other plasmas (**Figure 6A**), indicating that different epitopes are targeted by plasmas from the other donors.

**Fig. 6.**
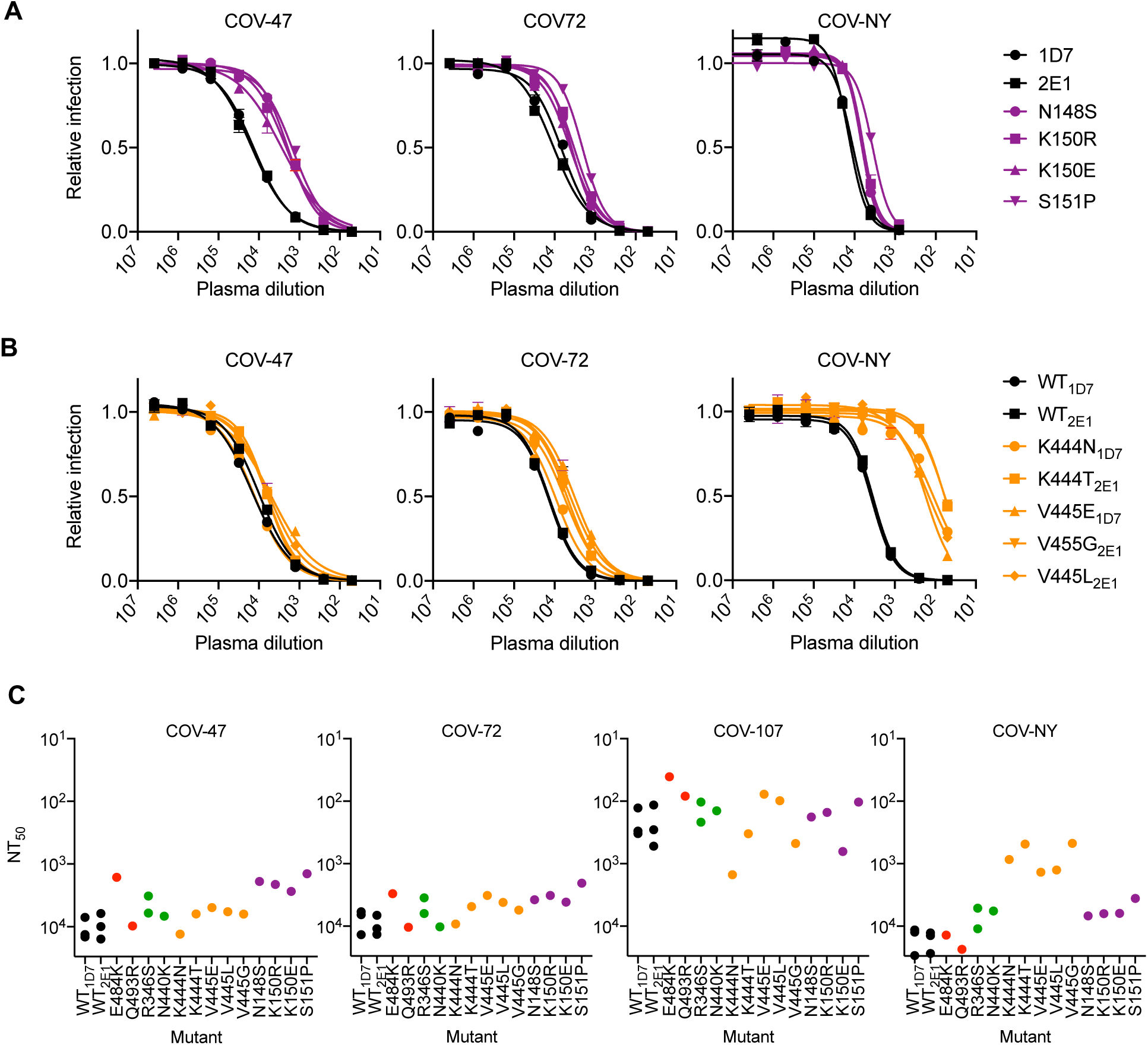
Neutralization of rVSV/SARS-CoV-2/GFP RBD mutants by convalescent plasma. A, B. Neutralization of rVSV/SARS-CoV-2/GFP mutants isolated following replication in the presence COV-47 plasma (A) or COV-NY plasma (B). 293T/ACE2cl.22 cells were inoculated with WT or mutant rVSV/SARS-CoV-2/GFP in the presence of increasing amounts of the indicated plasma, and infection quantified by flow cytometry, 16h later. C. Plasma neutralization sensitivity/resistance of rVSV/SARS-CoV-2/GFP mutants isolated following replication in the presence of monoclonal antibodies or convalescent plasma. NT_50_ values were calculated for each virus-plasma combination in two independent experiments.

The viral population that emerged during replication in COV-NY plasma yielded mutants K444N or T and V445G, E or L. Each of these mutations conferred substantial resistance to neutralization by COV-NY plasma, with ∼10-to 30-fold reduction in NT_50_ (**Figure 6B**). Thus, the dominant neutralizing activity in COV-NY plasma is represented by an antibody or antibodies recognizing an RBD epitope that includes K444 and V445. As was the case with COV-47 resistant mutants, viruses encoding the mutations conferring resistance to COV-NY plasma retained almost full sensitivity to neutralization by other plasmas (**Figure 6B**).

Interestingly, the mutations that conferred complete or near complete resistance to the potent RBD-specific monoclonal antibodies C121, C135 and C144 conferred little or no resistance to neutralization by plasma from the same individual, or other individuals (**Figure 6C**). These RBD-specific antibodies represent the most potent monoclonal antibodies isolated from COV-47, COV-72 and COV-107, respectively, but the retention of plasma sensitivity by the monoclonal antibody-resistant mutants suggests that these antibodies contribute little to the overall neutralization activity of plasma from the same individual. This finding is consistent with the observation that memory B cells producing these antibodies are rare (3), and with the results of the selection experiments in which rVSV/SARS-CoV-2/GFP replication in the presence of COV-47, COV-72 and COV-107 plasma did not enrich for mutations that correspond to the neutralization epitopes targeted by the monoclonal antibodies obtained from these individuals (**Figure 2 - supplement 1A**,**B, Figure 2 - supplement 2A**,**B. Table 2**). Overall, analysis of even this limited set of monoclonal antibodies and plasmas shows that potent neutralization can be conferred by antibodies that target diverse SARS-CoV-2 epitopes. Moreover, the most potently neutralizing antibodies generated in a given COVID19 convalescent individual may contribute in only a minor way to the overall neutralizing antibody response in that same individual.

### Natural occurrence of antibody-resistance RBD mutations

The aforementioned neutralizing antibody escape mutations were artificially generated during *in vitro* replication of a recombinant virus. However, as monoclonal antibodies are developed for therapeutic and prophylactic applications, and vaccine candidates are deployed, and the possibility of SARS-CoV-2 reinfection becomes greater, it is important both to understand pathways of antibody resistance and to monitor the prevalence of resistance conferring mutations in naturally circulating SARS-CoV-2 populations.

To survey the natural occurrence of mutations that might confer resistance to the monoclonal and plasma antibodies used in our experiments we used the GISAID (26) and CoV-Glue (27) SARS-CoV-2 databases. Among the 55,189 SARS-CoV-2 sequences in the CoV2-Glue database at the time of writing, 2175 different non-synonymous mutations were present in natural populations of SARS-CoV-2 S protein sequences. Consistent with the finding that none of the mutations that arose in our selection experiments gave an obvious fitness deficit (in the context of rVSV/SARS-CoV-2/GFP), most were also present in natural viral populations.

For phenotypic analysis of naturally occurring SARS-CoV-2 S mutations, we focused on the ACE2 interface of the RBD, as it is the target of most therapeutic antibodies entering clinical development (3, 17, 28), and is also the target of at least some antibodies present in convalescent plasmas. In addition to the mutations that arose in our antibody selection experiments, inspection of circulating RBD sequences revealed numerous naturally occurring mutations in the vicinity of the ACE2 binding site and the epitopes targeted by the antibodies (https://www.gisaid.org, http://cov-glue.cvr.gla.ac.uk) (**Figure 7A-D**). We tested nearly all of the mutations that are present in the GISAID database as of June 2020, in the proximity of the ACE2 binding site and neutralizing epitopes, for their ability to confer resistance to the monoclonal antibodies, using an HIV-1-based pseudotyped virus-based assay (**Figure 7A-C**). Consistent with, and extending our findings with rVSV/SARS-CoV-2/GFP, naturally occurring mutations at positions E484, F490, Q493, and S494 conferred complete or partial resistance to C121 and C144 (**Figure 7A, C**). While there was substantial overlap in the mutations that caused resistance to C121 and C144, there were also clear differences in the degree to which certain mutations (e.g. G446, L455R/I/F, F490S/L) affected sensitivity to the two antibodies. Naturally occurring mutations that conferred complete or partial resistance to C135 were at positions R346, N439, N440, K444, V445 and G446. In contrast to the C121/C144 epitope, these amino acids are peripheral to the ACE2 binding site on the RBD (**Figure 7D**). Indeed, in experiments where the binding of a conformationally stabilized trimeric S-NanoLuc fusion protein to 293T/ACE2cl.22 cells was measured, preincubation of S-NanoLuc with a molar excess of C121 or C144 completely blocked binding (**Figure 7E**). Conversely, preincubation with C135 only partly blocked binding to 293T/ACE2cl.22 cells, consistent with the finding that the C135 conformational epitope does not overlap the ACE2 binding site (**Figure 7D**). C135 might inhibit S-ACE-2 binding by steric interference with access to the ACE 2 binding site. These results are also consistent with experiments which indicated that C135 does not compete with C121 and C144 for binding to the RBD (3).

**Fig. 7.**
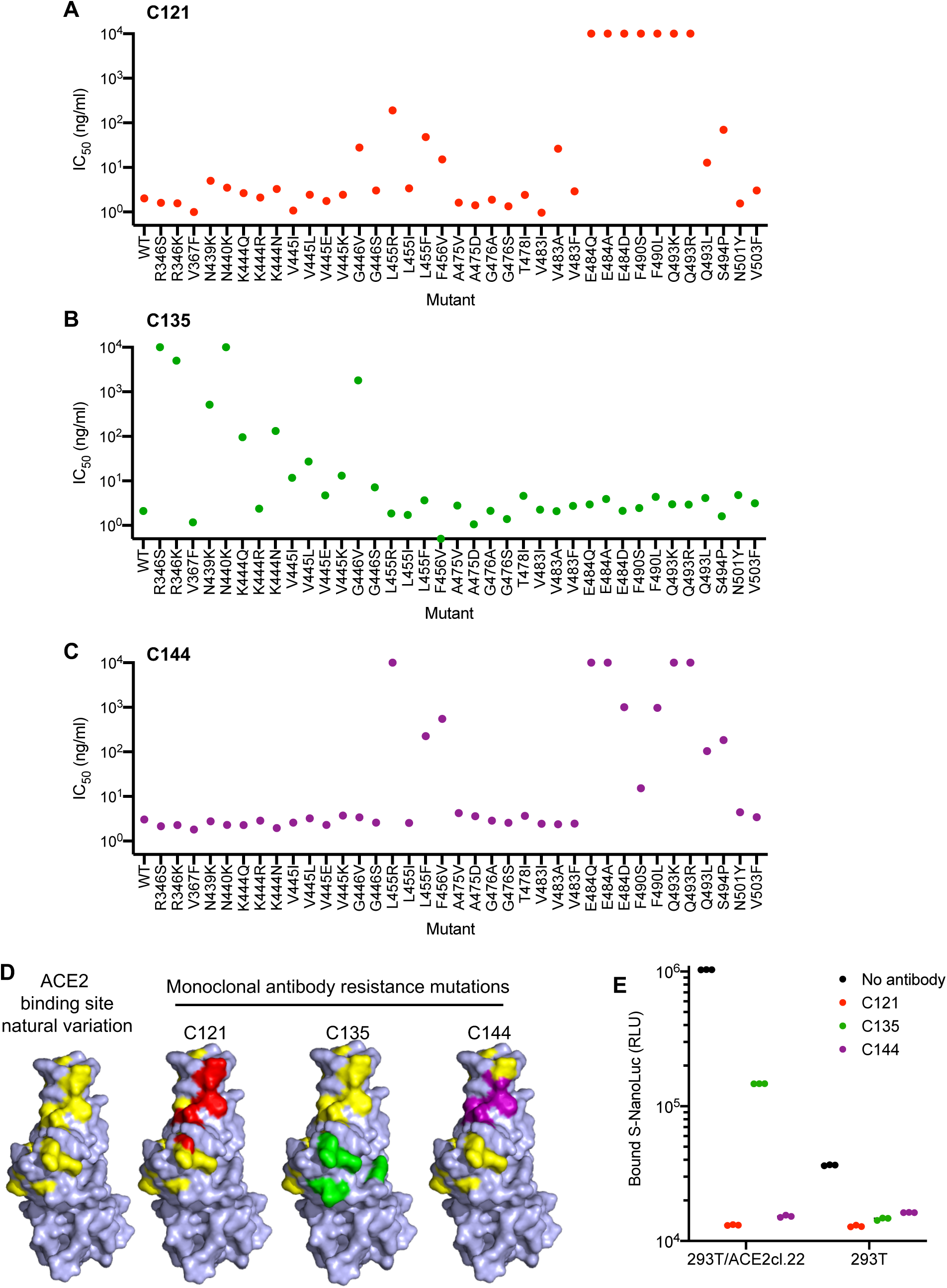
Effects of naturally occurring RBD amino acid substitutions on S sensitivity to neutralizing monoclonal antibodies. A-C. Neutralization of HIV-based reporter viruses pseudotyped with SARS-CoV-2 S proteins harboring the indicated naturally occurring substitutions. 293T/ACE2cl.22 cells were inoculated with equivalent doses of each pseudotyped virus in the presence of increasing amount of C121 (A) C132 (B) or C144 (C). IC_50_ values were calculated for each virus-antibody combination from two independent experiments. D. Position of substitutions conferring neutralization resistance relative to the amino acids close to the ACE2 binding site whose identity varies in global SARS-CoV-2 sequences. The RBD structure (from PDB 6M17 (32)) is depicted with naturally varying amino acids close to the ACE2 binding site colored in yellow. Amino acids whose substitution confers partial or complete (IC50 >10μg/ml) resistance to each monoclonal antibody in the HIV-pseudotype assays are indicated for C121 (red) C135 (green) and C144 (purple). E. Binding of S-NanoLuc fusion protein in relative light units (RLU) to 293T or 293T/ACE2cl.22 cells after preincubation in the absence or presence of C121, C135 and C144 monoclonal antibodies. Each symbol represents a technical replicate.

All of the mutations that were selected in our rVSV/SARS-CoV-2/GFP antibody selection experiments as well as other mutations that confer resistance to C121, C144 or C135 are found in naturally circulating SARS-CoV-2 populations at very low frequencies (**Figure 8**). With one exception (N439K) that is circulating nearly exclusively in Scotland and is present in ∼1% of COV-Glue database sequences, (and whose frequency may be overestimated due to regional oversampling) all antibody resistance mutations uncovered herein are present in global SARS-CoV-2 at frequencies of <1 in 1000 sequences (**Figure 8**). The frequency with which the resistance mutations are present in naturally occurring SARS-CoV-2 sequences appeared rather typical compared to other S mutations, with the caveat that sampling of global SARS-CoV-2 is nonrandom. Therefore, these observations do not provide evidence that the neutralizing activities exhibited by the monoclonal antibodies or plasma samples used herein have driven strong selection of naturally circulating SARS-CoV-2 sequences thus far (**Figure 8**).

**Fig. 8.**
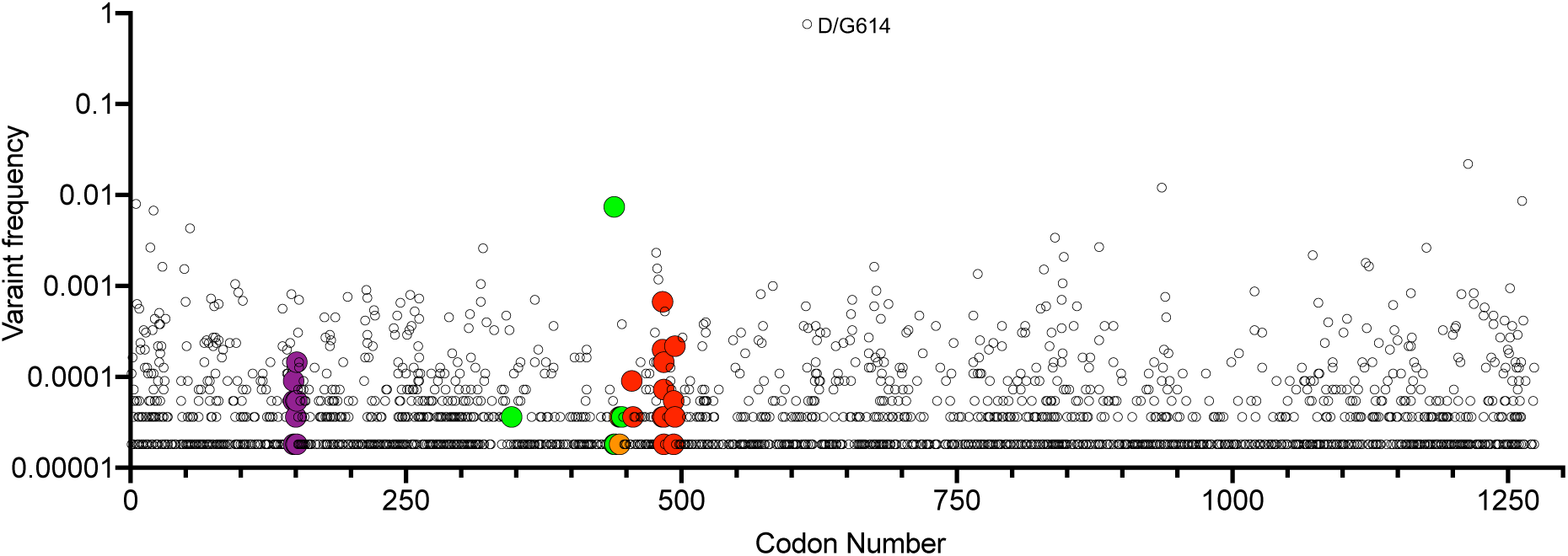
Position and frequency of S amino acid substitutions in SARS-CoV-2 S. Global variant frequency reported by CoV-Glue in the SARS-CoV-2 S protein. Each individual variant is indicated by a symbol whose position in the S sequence is indicated on the X-axis and frequency reported by COV-Glue is indicated on the Y-axis. Individual substitutions at positions where mutations conferring resistance to neutralizing antibodies or plasma were found herein are indicated by enlarged and colored symbols: red for C121 and C144, green for C135, purple for COV-47 plasma and orange for COV-NY plasma. The common D/G614 variant is indicated.

### Selection of combinations of monoclonal antibodies for therapeutic and prophylactic applications

The ability of SARS-CoV-2 monoclonal antibodies and plasma to select variants that are apparently fit and that naturally occur at low frequencies in circulating viral populations suggests that therapeutic use of single antibodies might select for escape mutants. To mitigate against the emergence or selection of escape mutations during therapy, or during population-based prophylaxis, we tested whether combinations of monoclonal antibodies could suppress the emergence of resistant variants during in vitro selection experiments. Specifically, we repeated antibody selection experiments in which rVSV/SARS-CoV-2/GFP populations containing 10^6^ infectious virions were incubated with 10μg/ml of each individual monoclonal antibody, or mixtures containing 5μg/ml of each of two antibodies (**Figure 9**). C121 and C144 target largely overlapping epitopes, and mutations conferring resistance to one of these antibodies generally conferred resistance to the other (**Figure 6**). Therefore, we used mixtures of antibodies targeting clearly distinct epitopes (C121+C135 and C144+C135). As previously, replication of rVSV/SARS-CoV-2/GFP in the presence of a single monoclonal antibody enabled the formation of infected foci in p1 cultures (**Figure 9A-C**), that rapidly expanded and enabled the emergence of apparently resistant virus populations. Indeed, rVSV/SARS-CoV-2/GFP yields from p2 cultures established with one antibody (C121, C135 or C144) were indistinguishable from those established with no antibody (**Figure 9D**). Conversely, rVSV/SARS-CoV-2/GFP replication in the presence of mixtures of C121+C135 or C144+C135 led to sparse infection of individual cells in p1 cultures, but there was little or no formation of foci that would suggest propagation of infection from these infected cells (**Figure 9A-C**), Therefore, it is likely that infected cells arose from rare, non-neutralized, virions that retained sensitivity to at least one of the antibodies in mixture. Consequently, viral spread was apparently completely suppressed and no replication competent rVSV/SARS-CoV-2/GFP was detected in p2 cultures established with mixtures of the two antibodies (**Figure 9D**).

**Fig. 9.**
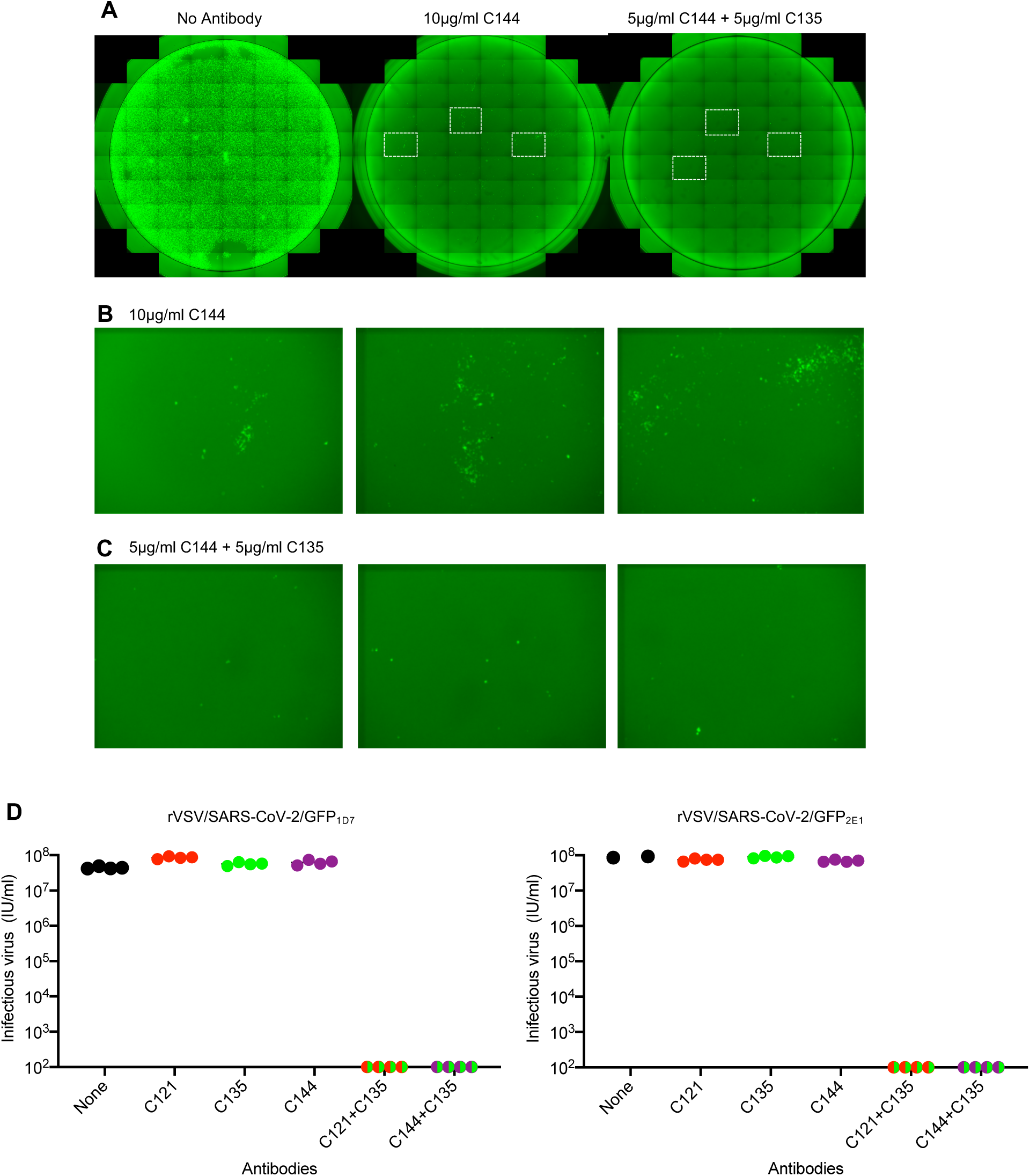
Suppression of antibody resistance through the use of antibody combinations. A. Representative images of 293T/ACE2(B) cells infected with the equivalent doses of rVSV/SARS-CoV-2/GFP in the absence or presence of 10μg/ml of one (C144) or 5μg/ml of each of two (C144 +C135) neutralizing monoclonal antibodies. B. Expanded view of the boxed areas containing individual plaques from the culture infected in the presence of 10μg/ml C144. C. Expanded view of the boxed areas in A containing infected cells from the culture infected in the presence of 5μg/ml each of C144 and C135). D. Infectious virus yield following two passages of rVSV/SARS-CoV-2/GFP in the absence or presence of individual neutralizing antibodies or combinations of two antibodies. Titers were determined on 293T/ACE2cl.22 cells. Each symbol represents a technical replicate and results from two independent experiments using rVSV/SARS-CoV-2/GFP_1D7_ and rVSV/SARS-CoV-2/GFP_2E1_ are shown.

## Discussion

The degree to which resistance will impact effectiveness of antibodies in SARS-CoV-2 therapeutic and vaccine settings is currently unclear (28). Notably, the inter-individual variation in SARS-CoV-2 sequences is low compared to many other RNA viruses, in part because coronaviruses encode a 3’-5’ exonuclease activity. The exonuclease activity provides a proofreading function the enhances replication fidelity and limits viral sequence diversification (29).

However, replication fidelity is but one of several variables that affect viral population diversity. One determinant of total viral diversity is population size. Many millions of individuals have been infected by SARS-CoV-2, and a single swab from an infected individual can contain in excess of 10^9^ copies of viral RNA (30). It follows that SARS-CoV-2 genomes encoding every possible single amino-acid substitution are present in the global population, and perhaps in a significant fraction of individual COVID19 patients. Thus, the frequency with which particular variants occur in the global SARS-CoV-2 population is strongly influenced by the frequency with which negative and positive selection pressures that favor their propagation are encountered, as well as founder effects at the individual patient and population levels.

Fitness effects of mutations will obviously vary and will suppress the prevalence of deleterious mutations. However, otherwise neutral, or even modestly deleterious, mutations will rise in prevalence if they confer escape from selective pressures, such as immune responses. Notably, the neutralizing antibody escape mutations described herein did not detectably alter rVSV/SARS-CoV-2/GFP replication, did not affect ACE2-Fc sensitivity and were found in natural populations at unexceptional frequencies, consistent with the notion that they do not have large effects on fitness in the absence of neutralizing antibodies. The prevalence of neutralizing antibody escape mutations will also be strongly influenced by the frequency with which SARS-CoV-2 encounters neutralizing antibodies. Peak viral burden in swabs and sputum, which likely corresponds to peak infectiousness and frequency of transmission events, appears to approximately correspond with the onset of symptoms, and clearly occurs before seroconversion (30). Thus, it is quite likely that most transmission events involve virus populations that are yet to experience antibody-imposed selective pressure in the transmitting individual. Such a scenario would reduce the occurrence of antibody escape mutations in natural viral populations. It will be interesting to determine whether viral sequences obtained late in infection are more diverse or have evidence of immunological escape mutations.

There are situations that are anticipated to increase the frequency of encounters between SARS-CoV-2 and antibodies that could impact the emergence of antibody resistance. Millions of individuals have already been infected with SARS-CoV-2 and among them, neutralizing antibody titers are extremely variable (3, 20, 21). It is plausible that those with weak immune responses or waning immunity can become re-infected, and if so, that encounters between SARS-CoV-2 and pre-existing but incompletely protective neutralizing antibodies might drive the selection of escape variants. In a similar manner, poorly immunogenic vaccine candidates, convalescent plasma therapy, and suboptimal monoclonal antibody treatment, particularly monotherapy(28), could create conditions to drive the acquisition of resistance to commonly occurring antibodies in circulating virus populations.

The extent to which SARS-CoV-2 evasion of individual antibody responses would have pervasive effects on the efficacy of vaccines and monoclonal antibody treatment/therapy will also be influenced by the diversity of neutralizing antibody responses within and between individuals. Analysis of potent neutralizing antibodies cloned by several groups indicates that potent neutralizing antibodies are commonly elicited, and very similar antibodies, such as those containing IGHV3-53 and IGHV3-66 can be found in different individuals (3, 18, 19). These findings imply a degree of homogeneity the among neutralizing antibodies that are generated in different individuals. Nevertheless, each of the four convalescent plasma tested herein had distinct neutralizing characteristics. In two of the four plasma tested, selection experiments suggested that a dominant antibody specificity was responsible for a significant fraction of the neutralizing capacity of the plasma. Nevertheless, the failure of single amino acid substitutions to confer complete resistance to any plasma strongly suggests the existence of multiple neutralizing specificities in each donor. The techniques described herein could be adapted to broadly survey the diversity of SARS-CoV-2 neutralizing specificities in many plasma samples following natural infection or vaccination, enabling a more complete picture of the diversity of SARS-CoV-2 neutralizing antibody responses to be developed. Indeed, the approach described herein can be used to map epitopes of potent neutralizing antibodies rapidly and precisely. It has an advantage over other epitope mapping approaches (such as array-based oligo-peptide scanning or random site-directed mutagenesis), in that selective pressure acts solely on the naturally formed, fusion-competent viral spike. While mutations outside the antibody binding sites might lead to resistance, the functional requirement will prohibit mutations that simply disrupt the native conformation. Indeed, we found that selective constraints caused escape variants to maintain replicative fitness.

Human monoclonal antibodies targeting both the NTD and RBD of SARS-CoV-2 have been isolated, with those targeting RBD being especially potent. As these antibodies are used clinically (17, 28), in therapeutic and prophylactic modes, it will be important to identify resistance mutations and monitor their prevalence in a way that is analogous to antiviral and antibiotic resistance monitoring in other infectious diseases. Moreover, as is shown herein, the selection of antibody mixtures with non-overlapping escape mutations should reduce the emergence of resistance and prolong the utility of antibody therapies in SARS-CoV-2 infection.

## Materials and Methods

### Plasmid constructs

A replication competent rVSV/SARS-CoV-2/GFP chimeric virus clone, encoding the SARS-CoV-2 spike protein lacking the C-terminal 18 codons in place of G, as well as GFP immediately upstream of the L (polymerase) has been previously described (24). The pHIV-1_NL_GagPol and pCCNG/nLuc constructs that were used to generate SARS-CoV-2 pseudotyped particles have been previously described (24). The pSARS-CoV-2 protein expression plasmid containing a C-terminally truncated SARS-CoV-2 S protein (pSARS-CoV-2_Δ19_) containing a synthetic human-codon optimized cDNA (Geneart) has been previously described (24) and was engineered to include BamHI, MfeI, BlpI and AgeI restriction enzyme sites flanking sequences encoding the RBD. Gibson assembly was used to introduce mutant RBD sequences into this plasmid, that were generated synthetically (g/eBlocks IDT) or by overlap extension PCR with primers that incorporated the relevant nucleotide substitutions.

### Cell lines

HEK-293T cells and derivatives were cultured in Dulbecco’s Modified Eagle Medium (DMEM) supplemented with 10% fetal bovine serum (FBS) at 37°C and 5% CO_2_. All cell lines have been tested negative for contamination with mycoplasma. Derivatives expressing ACE2 were generated by transducing 293T cells with CSIB(ACE2) vector and the uncloned bulk population 293T/ACE2(B) or a single-cell clone 293T/ACE2.cl22 (24) were used.

### Replication competent VSV/SARS-CoV-2/GFP chimeric virus

The generation of infectious rVSV/SARS-CoV-2/GFP chimeric viruses stocks has been previously described (24). Two plaque purified variants designated rVSV/SARS-CoV-2/GFP_1D7_ and rVSV/SARS-CoV-2/GFP_2E1_ that encode F157S/R685M (1D7) and D215G/R683G (2E1) substitutions were used in these studies.

### HIV-1/CCNanoLuc2AEGFP-SARS-CoV-2 pseudotype particles

The HIV-1/NanoLuc2AEGFP-SARS-CoV-2 pseudotyped virions were generated as previously described (24). Briefly, 293T cells were transfected with pHIV_NL_GagPol, pCCNanoLuc2AEGFP and a WT or mutant SARS-CoV-2 expression plasmid (pSARS-CoV-2_Δ19_) using polyethyleneimine. At 48h after transfection, the supernatant was harvested, clarified, filtered, aliquoted and stored at -80°C.

### Infectivity assays

To measure the infectivity of pseudotyped or chimeric viral particles, viral stocks were serially diluted and 100µl of each dilution added to 293T/ACE2cl.22 target cells plated at 1×10^4^ cells/well in 100 µl medium in 96-well plates the previous day. Cells were then cultured for 48h hours (HIV-1 pseudotyped viruses) or 16h (replication competent rVSV/SARS-CoV-2/GFP), unless otherwise indicated, and then photographed or harvested for NanoLuc luciferase or flow cytometry assays.

### Selection of viruses in the presence of antibodies

For selection of viruses resistant to plasma or monoclonal antibodies, rVSV/SARS-CoV-2/GFP_1D7_ and rVSV/SARS-CoV-2/GFP_2E1_ populations containing 10^6^ infectious particles were incubated with dilutions of monoclonal antibodies (10μg/ml, 5μg/ml) or COVID19 plasma (1:50, 1:250, 1:500) for 1h at 37°C. Then, the virus antibody mixtures were incubated with 2×10^5^ 293T/ACE2(B) cells in 12 well plates. Two days later, the cells were imaged and supernatant from the wells containing the highest concentration of plasma or monoclonal antibodies that showed evidence of viral replication (GFP positive foci) or large numbers of GFP positive cells was harvested. A 100μl of the cleared supernatant was incubated with the same dilution of plasma or monoclonal antibody and then used to infect 2×10^5^ 293T/ACE2(B) cells in 12 well plates, as before. rVSV/SARS-CoV-2/GFP_1D7_ and rVSV/SARS-CoV-2/GFP_2E1_ were passaged in the presence of C121 or C144 two times before complete escape was apparent. rVSV/SARS-CoV-2/GFP_1D7_ and rVSV/SARS-CoV-2/GFP_2E1_ were passaged with C135 or plasma samples up to 5 times.

To isolate individual mutant viruses, selected rVSV/SARS-CoV-2/GFP_1D7_ and rVSV/SARS-CoV-2/GFP_2E1_ populations were serially diluted in medium without antibodies and individual viral variants isolated by visualizing single GFP-positive plaques at limiting dilutions in 96-well plates containing 1×10^4^ 293T/ACE2(B) cells. These plaque purified viruses were expanded, and further characterized using sequencing, spreading replication and neutralization assays.

### Sequence analyses

For the identification of putative antibody resistance mutations, RNA was isolated from aliquots of supernatant containing selected viral populations or individual plaque purified variants using Trizol-LS. The purified RNA was subjected to reverse transcription using random hexamer primers and Superscript III reverse transcriptase (Thermo Fisher Scientific, US). The cDNA was amplified using Phusion (NEB, US) and primers flanking RBD encoding sequences. Alternatively, a fragment including the entire S-encoding sequence was amplified using primers targeting VSV-M and VSV-L. The PCR products were gel-purified and sequenced either using Sanger-sequencing or NGS as previously described (31). Briefly, 1 µl of diluted DNA was used for the tagmentation reactions with 0.25 µl Nextera TDE1 Tagment DNA enzyme (catalog no. 15027865), and 1.25 µl TD Tagment DNA buffer (catalog no. 15027866; Illumina).

Subsequently, the DNA was ligated to unique i5/i7 barcoded primer combinations using the Illumina Nextera XT Index Kit v2 and KAPA HiFi HotStart ReadyMix (2X; KAPA Biosystems) and purified using AmPure Beads XP (Agencourt), after which the samples were pooled into one library and subjected to paired-end sequencing using Illumina MiSeq Nano 300 V2 cycle kits (Illumina) at a concentration of 12pM.

For analysis of NGS data, the raw paired-end reads were pre-processed to remove adapter sequences and trim low-quality reads (Phred quality score <20) using BBDuk. Filtered reads were mapped to the codon-optimized SARS-CoV-2 S sequence in rVSV/SARS-CoV-2/GFP using Geneious Prime (Version 2020.1.2). Mutations were annotated using Geneious Prime, with a P-value cutoff of 10^−6^. Information regarding RBD-specific variant frequencies, their corresponding P-values, and read depth were compiled using the Python programming language (version 3.7) running pandas (1.0.5), numpy (1.18.5), and matplotlib (3.2.2).

### Neutralization assays

To measure neutralizing antibody activity in plasma, serial dilutions of plasma beginning with a 1:12.5 or a 1:100 (for plasma COV-NY) initial dilution were five-fold serially diluted in 96-well plates over six or eight dilutions. For monoclonal antibodies, the initial dilution started at 40µg/ml. Thereafter, approximately 5×10^4^ infectious units of rVSV/SARS-CoV-2/GFP or 5×10^3^ infectious units of HIV/CCNG/nLuc/SARS-CoV-2 were mixed with the plasma or mAb at a 1:1 ratio and incubated for 1 hour at 37°C in a 96-well plate. The mixture was then added to 293T/ACE2cl.22 target cells plated at 1×10^4^ cells/well in 100 µl medium in 96-well plates the previous day. Thus, the final starting dilutions were 1:50 or 1:400 (for COV-NY) for plasma and 10µg/ml for monoclonal antibodies. Cells were then cultured for 16h (for rVSV/SARS-CoV-2/GFP) or 48h (for HIV/CCNG/nLuc/SARS-CoV-2). Thereafter, cells were harvested for flow cytometry or NanoLuc luciferase assays.

### Antibody binding and ACE2 binding inhibition assay

A conformationally stabilized (6P) version of the SARS-CoV-2 S protein(25), appended at its C-terminus with a trimerization domain, a GGSGGn spacer sequence, NanoLuc luciferase, Strep-tag, HRV 3C protease cleavage site and 8XHis (S-6P-NanoLuc) was expressed and purified from the supernatant of 293T Expi cells. Mutants thereof were also expressed and purifies following substitution of sequences encoding the RBD that originated from the unmodified S-expression plasmids

For antibody binding assays, 20ng, 40ng, or 80ng S-6P-NanoLuc (or mutants thereof) were mixed with 100ng of antibodies, C121, C135, or C144, \ diluted in LI-COR Intercept blocking buffer, in a total volume of 60μl/well in 96-well plate. After a 30 min incubation, 10ul protein G magnetic beads was added to each well and incubated for 1.5 hr. The beads were then washed 3 times and incubated with 30ul lysis buffer (Promega). Then 15μl of the lysate was used to measure bound NanoLuc activity.

For ACE2-binding inhibition assays, 20ng of S-6P-NanoLuc was mixed with 100ng of antibodies, C121, C135 or C144, diluted in 3% goat serum/PBS, in a total volume of 50μl. After 30 minutes incubation, the mixture was incubated with 1×10^5^ 293T cells, or 293T/ACE2cl.22 cells for 2 hours at 4°C. The cells were then washed 3 times and lysed with 30μl lysis buffer and 15μl of the lysate was used to measure bound NanoLuc activity.

### Reporter gene assays

For the NanoLuc luciferase assays, cells were washed gently, twice with PBS and lysed in Lucifersase Cell culture Lysis reagent (Promega). NanoLuc luciferase activity in the lysates was measured using the Nano-Glo Luciferase Assay System (Promega) and a Modulus II Microplate Multimode reader (Turner BioSystem) or a Glowmax Navigator luminometer (Promega), as described previously(24). To record GFP+ cells, 12-well plates were photographed using an EVOS M7000 automated microscope. For flow cytometry, cells were trypsinized, fixed and enumerated using an Attune NxT flow cytometer. The half maximal inhibitory concentrations for plasma (NT_50_), and monoclonal antibodies (IC_50_) was calculated using 4-parameter nonlinear regression curve fit to raw or normalized infectivity data (GraphPad Prism). Top values were unconstrained, the bottom values were set to zero.

### Human plasma samples and monoclonal antibodies

The human plasma samples COV-47, COV-72 and COV-107 and monoclonal antibodies C144, C135 and C121 used in this study were previously reported (3). The human plasma sample COV-NY was obtained from the New York Blood Center(21). All plasma samples were obtained under protocols approved by Institutional Review Boards at both institutions.

## Author Contributions

PDB, TH, YW, FS, MCN, CMR and DR conceived and supervised the studies. FS and YW constructed all recombinant plasmids, viruses and cell lines and did all the experiments with the following exceptions: FZ did the ACE2 binding assay, DP and JCCL assisted with illumina sequencing and analysis, TH and JDS did the HIV-1 pseudotype neutralization assays. FM, FS and JCCL developed the HIV-1 pseudovirus assay. MR provided general technical assistance. MC, CG, LL and CDH provided clinical samples, MA, AC and ZW cloned monoclonal antibodies that were produced and purified by AG and MC. PDB wrote the manuscript with input from other authors.

## Acknowledgements

We acknowledge the generous contributions the GISAID database contributors and curators. This work was supported by NIH grants P01AI138398-S1, 2U19AI111825 (to M.C.N. and C.M.R), R01AI091707-10S1 (to C.M.R.) R01AI078788 (to T.H.) R37AI64003 (to P.D.B.) and by the George Mason University Fast Grant (to D.F.R. and to C.M.R.) and the European ATAC consortium EC101003650 (to D.F.R) and the G. Harold and Leila Y. Mathers Charitable Foundation (to C.M.R.). C.G. was supported by the Robert S. Wennett Post-Doctoral Fellowship, in part by the National Center for Advancing Translational Sciences (National Institutes of Health Clinical and Translational Science Award program, grant UL1 TR001866), and by the Shapiro-Silverberg Fund for the Advancement of Translational Research. P.D.B. and M.C.N. are Howard Hughes Medical Institute Investigators.

---

**Figure 2 - supplement 1.**
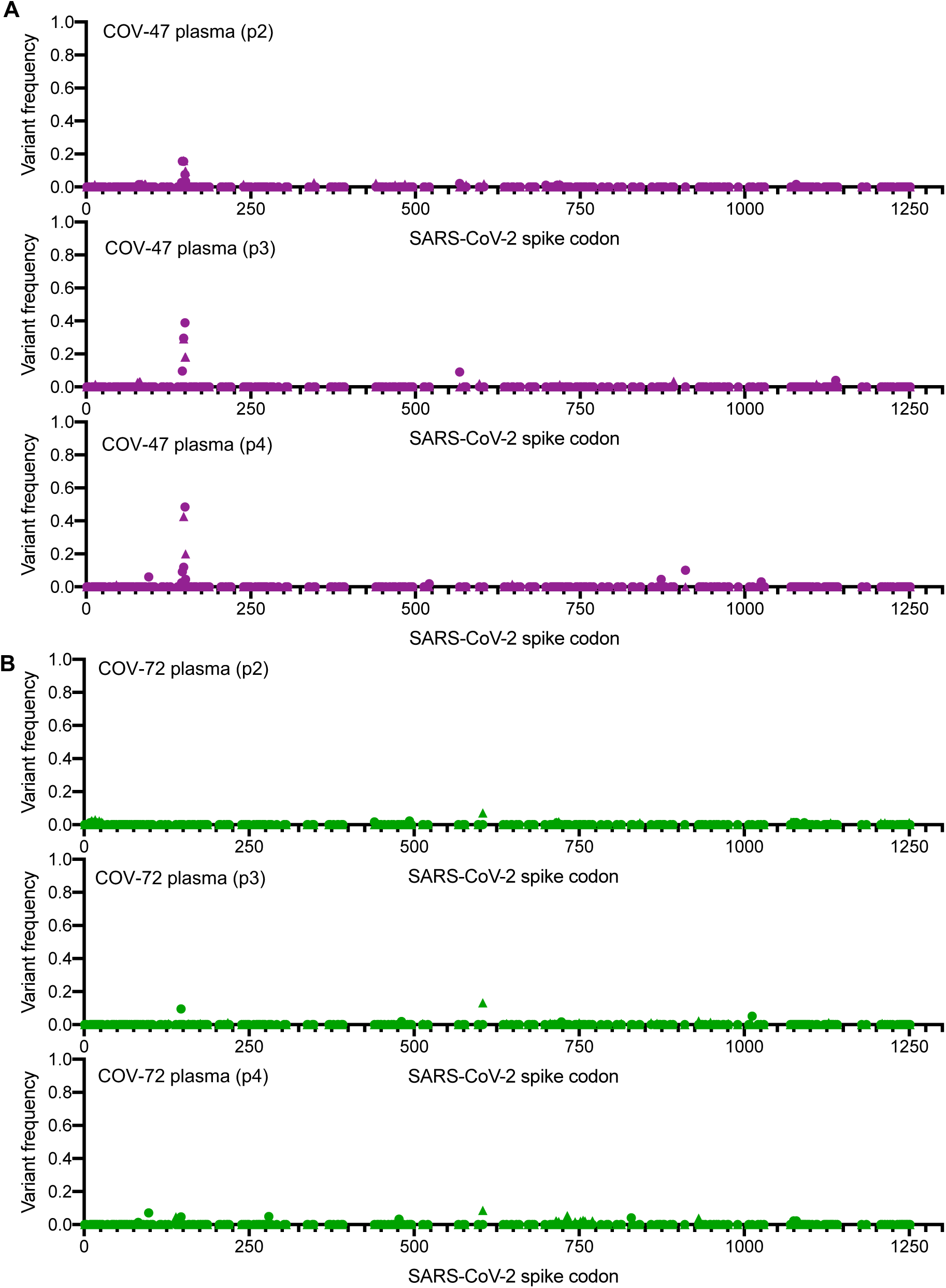
Analysis of S-encoding sequences following rVSV/SARS-CoV-2/GFP replication in the presence of convalescent plasma COV-47 and COV-72. A-B. Graphs depict the S codon position (X-axis) and the frequency of non-synonymous substitutions (Y-axis) following the second, third or fourth passage (p2-p4) of rVSV/SARS-CoV-2/GFP on 293T/ACE2(B) cells in the presence of COV-47 plasma (A), or COV-72 plasma (B). Results are shown for both rVSV/SARS-CoV-2/GFP variants (the frequency of 1D7 mutations is plotted as circles and 2E1 mutations as triangles).

**Figure 2 - supplement 2.**
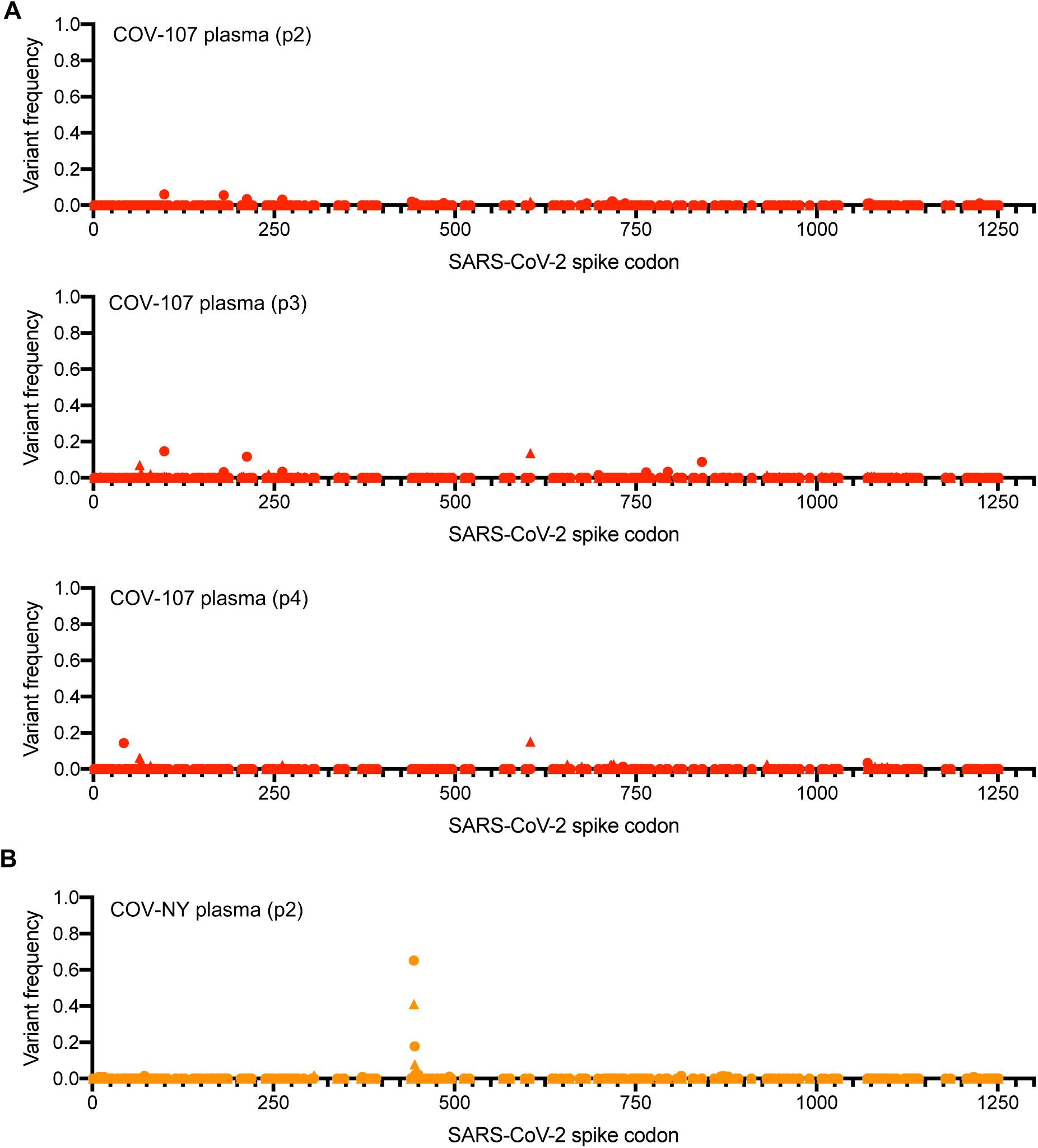
Analysis of S-encoding sequences following rVSV/SARS-CoV-2/GFP replication in the presence of convalescent plasma COV-107 and COV-NY. A-B. Graphs depict the S codon position (X-axis) and the frequency of non-synonymous substitutions (Y-axis) following the second, third or fourth passage (p2-p4) of rVSV/SARS-CoV-2/GFP on 293T/ACE2(B) cells in the presence of COV-107 plasma (A), or second passage in the presence of COV-NY plasma (B). Results are shown for both rVSV/SARS-CoV-2/GFP variants (the frequency of 1D7 mutations is plotted as circles and 2E1 mutations as triangles).

**Figure 3 - supplement 1.**
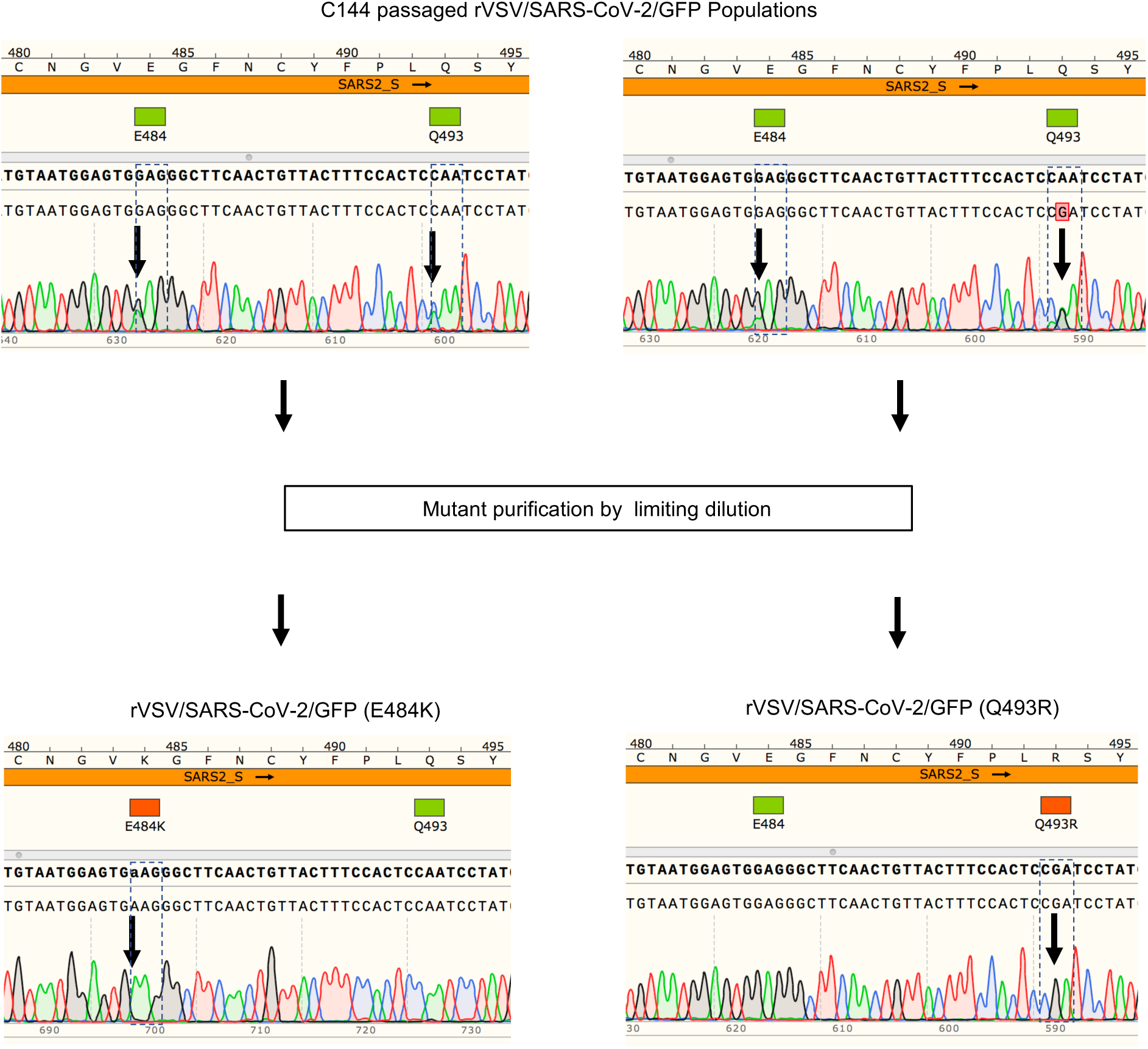
Example of plaque purification of individual viral mutants from populations passaged in the presence of antibodies. The upper panels show sequence traces from amplicons obtained from viral populations following replication in the presence of monoclonal antibodies, the bottom panels show sequence traces of amplicons obtained from mutants isolated by limiting dilution of the viral populations

## References

1. Plotkin SA. Correlates of Protection Induced by Vaccination. Clinical and Vaccine Immunology. 2010;17:1055–65.

2. Kellam P, Barclay W. The dynamics of humoral immune responses following SARS-CoV-2 infection and the potential for reinfection. The Journal of general virology. 2020.

3. Robbiani DF, Gaebler C, Muecksch F, Lorenzi JCC, Wang Z, Cho A, et al. Convergent Antibody Responses to SARS-CoV-2 Infection in Convalescent Individuals. Nature. 2020;In press.

4. Brouwer PJM, Caniels TG, van der Straten K, Snitselaar JL, Aldon Y, Bangaru S, et al. Potent neutralizing antibodies from COVID-19 patients define multiple targets of vulnerability. Science. 2020;In Press.

5. Cao Y, Su B, Guo X, Sun W, Deng Y, Bao L, et al. Potent neutralizing antibodies against SARS-CoV-2 identified by high-throughput single-cell sequencing of convalescent patients’ B cells. Cell. 2020;182:1–12.

6. Chen X, Li R, Pan Z, Qian C, Yang Y, You R, et al. Human monoclonal antibodies block the binding of SARS-CoV-2 spike protein to angiotensin converting enzyme 2 receptor. Cellular & molecular immunology. 2020;17(6):647–9.

7. Chi X, Yan R, Zhang J, Zhang G, Zhang Y, Hao M, et al. A potent neutralizing human antibody reveals the N-terminal domain of the Spike protein of SARS-CoV-2 as a site of vulnerability. bioRxiv. 2020.

8. Ju B, Zhang Q, Ge X, Wang R, Yu J, Shan S, et al. Potent human neutralizing antibodies elicited by SARS-CoV-2 infection. Nature. 2020;In press.

9. Rogers TF, Zhao F, Huang D, Beutler N, Burns A, He W-t, et al. Isolation of potent SARS-CoV-2 neutralizing antibodies and protection from disease in a small animal model. Science. 2020;In press.

10. Shi R, Shan C, Duan X, Chen Z, Liu P, Song J, et al. A human neutralizing antibody targets the receptor binding site of SARS-CoV-2. Nature. 2020;In press.

11. Wu Y, Wang F, Shen C, Peng W, Li D, Zhao C, et al. A non-competing pair of human neutralizing antibodies block COVID-19 virus binding to its receptor ACE2. medRxiv. 2020.

12. Zost SJ, Gilchuk P, Case JB, Binshtein E, Chen RE, Reidy JX, et al. Potently neutralizing human antibodies that block SARS-CoV-2 receptor binding and protect animals. bioRxiv. 2020.

13. Wec AZ, Wrapp D, Herbert AS, Maurer D, Haslwanter D, Sakharkar M, et al. Broad Neutralization of SARS-related Viruses by Human Monoclonal Antibodies. Science. 2020;In press.

14. Seydoux E, Homad LJ, MacCamy AJ, Parks KR, Hurlburt NK, Jennewein MF, et al. Characterization of neutralizing antibodies from a SARS-CoV-2 infected individual. bioRxiv. 2020.

15. Liu L, Wang P, Nair MS, Yu J, Rapp M, Wang Q, et al. Potent Neutralizing Antibodies Directed to Multiple Epitopes on SARS-CoV-2 Spike. bioRxiv. 2020.

16. Kreer C, Zehner M, Weber T, Ercanoglu MS, Gieselmann L, Rohde C, et al. Longitudinal Isolation of Potent Near-Germline SARS-CoV-2-Neutralizing Antibodies from COVID-19 Patients. Cell. 2020.

17. Hansen J, Baum A, Pascal KE, Russo V, Giordano S, Wloga E, et al. Studies in humanized mice and convalescent humans yield a SARS-CoV-2 antibody cocktail. Science. 2020.

18. Barnes CO, West AP, Jr., Huey-Tubman KE, Hoffmann MAG, Sharaf NG, Hoffman PR, et al. Structures of Human Antibodies Bound to SARS-CoV-2 Spike Reveal Common Epitopes and Recurrent Features of Antibodies. Cell. 2020.

19. Yuan M, Liu H, Wu NC, Lee CD, Zhu X, Zhao F, et al. Structural basis of a shared antibody response to SARS-CoV-2. Science. 2020.

20. Wu F, Wang A, Liu M, Wang Q, Chen J, Xia S, et al. Neutralizing antibody responses to SARS-CoV-2 in a COVID-19 recovered patient cohort and their implications. medRxiv. 2020.

21. Luchsinger LL, Ransegnola B, Jin D, Muecksch F, Weisblum Y, Bao W, et al. Serological Analysis of New York City COVID19 Convalescent Plasma Donors. medRxiv. 2020.

22. Bloch EM, Shoham S, Casadevall A, Sachais BS, Shaz B, Winters JL, et al. Deployment of convalescent plasma for the prevention and treatment of COVID-19. The Journal of clinical investigation. 2020;130(6):2757–65.

23. Al-Riyami AZ, Schäfer R, van der Berg K, Bloch EM, Escourt LJ, Goel R, et al. Clinical use of Convalescent Plasma in the COVID-19 pandemic; a transfusion-focussed gap analysis with recommendations for future research priorities. Vox sanguinis. 2020.

24. Schmidt F, Weisblum Y, Muecksch F, Hoffmann HH, Michailidis E, Lorenzi JCC, et al. Measuring SARS-CoV-2 neutralizing antibody activity using pseudotyped and chimeric viruses. The Journal of experimental medicine. 2020;In press.

25. Hsieh CL, Goldsmith JA, Schaub JM, DiVenere AM, Kuo HC, Javanmardi K, et al. Structure-based Design of Prefusion-stabilized SARS-CoV-2 Spikes. bioRxiv. 2020.

26. Elbe S, Buckland-Merrett G. Data, disease and diplomacy: GISAID’s innovative contribution to global health. Global challenges (Hoboken, NJ). 2017;1(1):33–46.

27. Singer J, Gifford R, Cotten M, Robertson D. CoV-GLUE: A Web Application for Tracking SARS-CoV-2 Genomic Variation. Preprints. 2020.

28. Baum A, Fulton BO, Wloga E, Copin R, Pascal KE, Russo V, et al. Antibody cocktail to SARS-CoV-2 spike protein prevents rapid mutational escape seen with individual antibodies. Science. 2020.

29. Denison MR, Graham RL, Donaldson EF, Eckerle LD, Baric RS. Coronaviruses: an RNA proofreading machine regulates replication fidelity and diversity. RNA biology. 2011;8(2):270–9.

30. Wölfel R, Corman VM, Guggemos W, Seilmaier M, Zange S, Müller MA, et al. Virological assessment of hospitalized patients with COVID-2019. Nature. 2020;581(7809):465–9.

31. Gaebler C, Lorenzi JCC, Oliveira TY, Nogueira L, Ramos V, Lu CL, et al. Combination of quadruplex qPCR and next-generation sequencing for qualitative and quantitative analysis of the HIV-1 latent reservoir. The Journal of experimental medicine. 2019;216(10):2253–64.

32. Yan R, Zhang Y, Li Y, Xia L, Guo Y, Zhou Q. Structural basis for the recognition of SARS-CoV-2 by full-length human ACE2. Science. 2020;367(6485):1444–8.

